# Toll-like receptor 9 contributes in microglial activation and lysosomal dysfunction to promote Alzheimer’s disease

**DOI:** 10.64898/2026.01.16.699962

**Authors:** Moïse de Lavaissière de Lavergne, Lucie Mazzola, Katrina Podsypanina, Cecilia Estrella, Philippe Verwaerde, Irini Evnouchidou, Guillaume Darrasse-Jèze, Marie-Claude Potier, Sheela Vyas, Bénédicte Manoury

**Affiliations:** Institut Necker Enfants Malades, INSERM U1151-CNRS UMR 8253, Université Paris Cité, Faculté de Médecine Necker, Paris, France; Alzprotect, Therapeutics solution for neurodegeneration, Loos, France; Centre de Recherche sur l’Inflammation, U1149 INSERM, CNRS ERL8252, Université de Paris Cité, Faculté de Médecine Bichat, Paris, France; Sorbonne Université, Institut du Cerveau - Paris Brain Institute - ICM, CNRS, APHP, Hôpital de La Pitié Salpêtrière, INSERM, Paris, France; Laboratory of Gene Regulation and Adaptive Behaviors, Department of Neuroscience Paris Seine, INSERM U1130, CNRS UMR 8246, UPMC UM 119, Université Pierre et Marie Curie, Paris, France

## Abstract

Toll like receptor 9 (TLR9) is an endosomal-lysosomal innate immune receptor which recognizes microbial and host-derived double-stranded DNA (dsDNA) to induce cellular immunity. TLR9 pathway has been implicated in Alzheimer’s disease (AD) pathology via its ability to modulate peripheral immune cells but its exact role in the central nervous system (CNS) remains unclear. Here we show that extranuclear dsDNA accumulating in human AD brain and in a mouse model of AD, together with microglia around amyloid-β (Aβ) plaques stimulates TLR9 *in vitro*. Loss of TLR9 inhibits the transition of homeostatic microglia to reactive microglia, a cell type associated with a chronic pro-inflammatory state, promoting their reprogramming into an anti-inflammatory state, thereby restoring lysosomal integrity required for Aβ peptides degradation. Predisposition to lysosome permeabilization could be mimicked with repeated TLR9 stimulation using CpG-A ODNs in a human microglial cell line and was rescued with TLR9 inhibition. Furthermore, lack of TLR9 protects mice developing AD from cognitive defects and neuronal loss. Aβ pathology is also reduced specifically in male mice. Altogether, our results have identified a critical role of TLR9 in AD, where it modulates microglial activation and lysosome function to interfere with the progression of AD. These findings suggest that targeting TLR9 signaling and lysosomal function may be a relevant strategic approach to reduce AD pathology.

## Introduction

AD is a neurodegenerative disease which is characterized by amyloid-β (Aβ) plaque deposition, neurofibrillary tangle formation due to tau hyperphosphorylation, and associated with neurodegeneration and cognitive defects. Genetic studies in AD reveal a fundamental role of innate immune responses in neurodegenerative processes stemming from Aβ and tau pathologies. As these responses either counteract or exacerbate AD pathology, understanding their actions is essential. Toll-like receptors (TLRs), major initiators of immune activation, are potentially decisive in modulating AD pathology, yet their exact roles in disease remain unclear. In this respect, involvement of TLR2 and TLR4 in Aβ-mediated activation of microglia or in Aβ clearance capacity has been reported (Reed-Gheagan et al. 2009, McDonald et al. 2016, Fujikura et al. 2019). Multiple injections of LPS, a TLR4 ligand, in the periphery were also shown to lead to long---term microglial ‘memory’ that modifies Aβ pathology through epigenetic regulation of microglial inflammatory and clearance function (Wendeln et al. 2018).

TLR9 is an endosomal-lysosomal receptor stimulated upon recognition of CpG sequences found in DNA from pathogens but also in mitochondrial DNA (mtDNA) released from dying neurons. TLR9 is particularly interesting in AD: its activation by peripheral injections of its ligand CpG leads to a reduction in both Aβ and tau pathologies in several models of AD (Scholtzova et al. 2014, Patel et al. 2021). TLR9 signalling is regulated by the exonuclease phospholipase D3 (PLD3) and the glycoprotein progranulin (PGRN), whose genes are linked to AD predisposition (Lee et al. 2011, Cruchaga et al. 2013). PLD3 inhibits TLR9 signalling by breaking down TLR9 ligands in the lysosome thereby limiting their availability for stimulation. Thus, immune cells lacking PLD3 show massive inflammatory response due to exacerbated TLR9 activation (Gavin et al. 2018). Granulin (GRN) is a soluble factor which potentiates TLR9 signalling by binding to TLR9 ligand and facilitates its delivery in the lysosomal compartment, as absence of GRN leads to reduced TLR9 response (Park et al. 2011). Furthermore, the beneficial effect of AZP2006 (a compound in phase 2 clinical trial for progressive supranuclear palsy, a tauopathy) was reported to occur through stabilization of PGRN, which also inhibited TLR9 signaling (Callizot et al. 2021). Low levels of cell-free mtDNA were reported in cerebrospinal fluid (CSF) of AD patients (Podlesniy et al. 2013) which could potentially activate TLR9 in immune cells including microglia as it was recently shown (Pinti et al. 2021). Moreover, TLR9 is activated by asparagine endopeptidase (AEP) cleavage in lysosomes (Sepulveda et al. 2009). AEP can also cleave amyloid precursor protein (APP) to produce Aβ, and its cleavage of tau results in its hyperphosphorylation (Basurto-Islas et al. 2013, Zhang et al. 2015). Lysosome fitness in microglia is critical for the clearance of protein aggregates and neuronal debris through phagocytosis. However, the interaction between TLR9 trafficking from the endoplasmic reticulum (ER) followed by its processing in lysosomes in relation to pathological protein aggregate clearance remains to be determined. While these results indicate that TLR9 may be involved, it is not known how TLR9 activation affects AD physiopathology.

We hypothesized that chronic TLR9 activation by double-stranded DNA (dsDNA) release leads to microglial dysfunction and promotes AD progression. In this study, we found an increase in the expression of extranuclear dsDNA in the brain of human AD patients and of mice developing AD. The absence of TLR9 reduced Aβ pathology, restored cognitive deficits and induced an anti-inflammatory microglial phenotype that contributed to limit neuroinflammation and restore lysosomal functions for effective Aβ deposit clearance. These findings provide a new line of evidence that the innate immune receptor TLR9 plays a critical role in AD pathology.

## Results

### dsDNA release and activation of TLR9 pathway during AD

Accumulation of DNA double stranded breaks (DSB) and release of mtDNA due to defective mitophagy lead to the generation of TLR9 ligands and are associated with ageing and the development of AD (Jylhävä et al. 2011, Simpson et al. 2015). To assess the role of TLR9 in AD pathology, we crossed TLR9-deficient mice with APP23 mice (A9KO), a model for familial AD. APP23 mice have a sevenfold overexpression of the human amyloid precursor protein (APP) with the Swedish AD-linked double mutation (K670N/M671L) under the control of the Thy1 neuron promoter and are associated with progressive Aβ deposition. APP23 mice start to develop plaques and inflammatory reactions by 7-8 months of age in the cortex and show memory-related learning impairments. To verify the presence of potential TLR9 ligands in the CNS of these mice, we immuno-stained two regions of the brain, the hippocampus (CA3 region) and the entorhinal cortex where Aβ plaque deposition occurs, for the presence of dsDNA. We found a strong accumulation of extranuclear dsDNA, in the brain of 12 months and 18 months old APP23 mice in comparison to A9KO and WT mice (Figure 1A-B). Furthermore, for all genotypes, this extranuclear dsDNA was mostly localized outside of mitochondrial and PGRN positive compartments (Figure S1A-D). Pathology was also associated with increased mitochondrial mass at early stages of Aβ deposition, especially in the hippocampus as evidenced by Tom20 immunostaining (Figure S1E-F) and mtDNA purified from APP23 mouse brain (Figure S1G) stimulated TLR9 *in vitro* (Figure 1C). Additionally, dsDNA in the brain of APP23 mice was detected in the vicinity of dense core plaques stained by thioflavin S (ThioS) and co-localized with recruited microglia (Figure 1D). Microglia from healthy mice also express TLR9 in both of these regions (Figure S1H) and *in vitro* TLR9 activation of microglia purified from APP23 mice led to increased production of TNF-α and IL-6 compared to WT microglia (Figure S1I).

**Figure 1.**
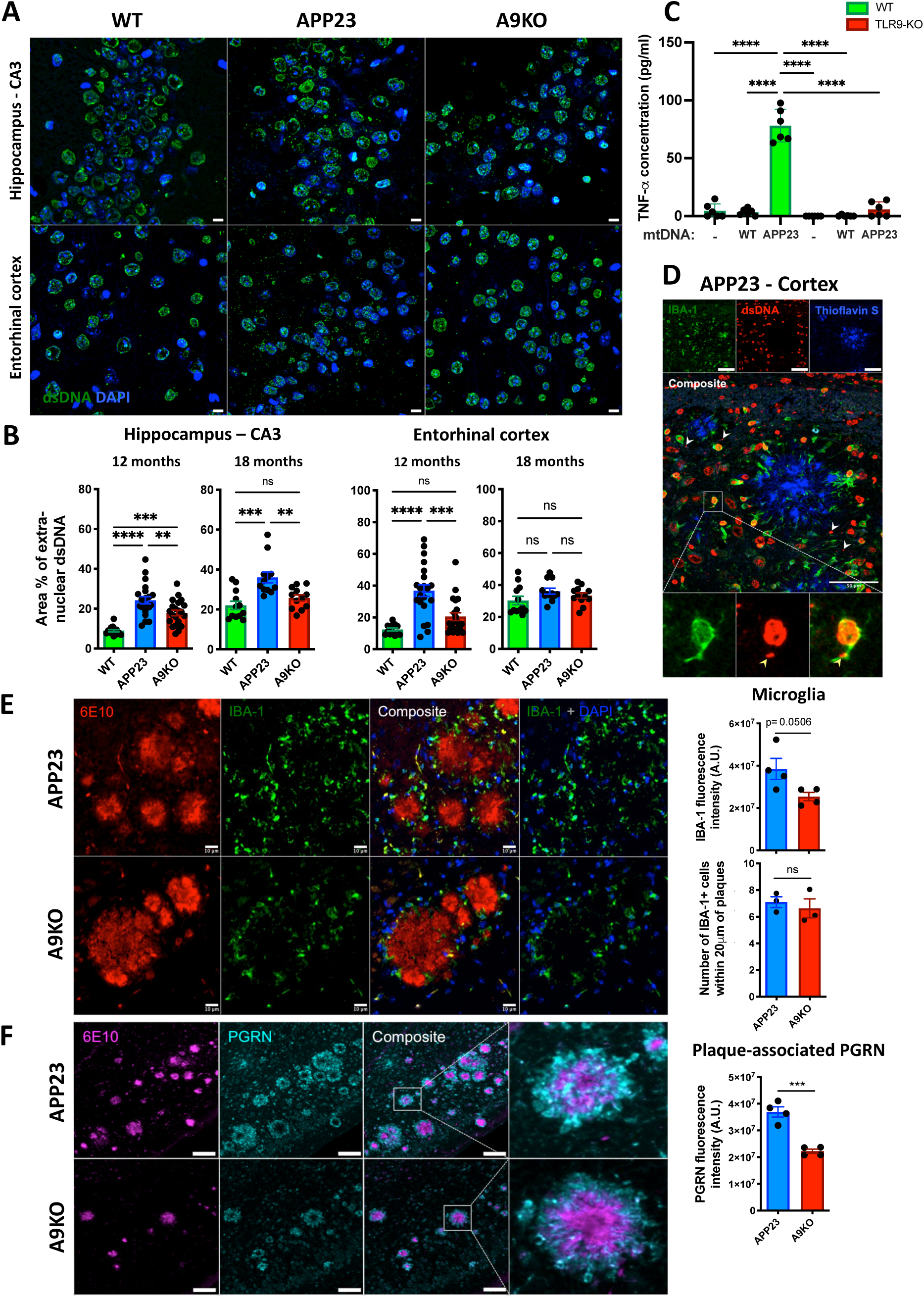
dsDNA release and microglia activation during AD. **(A)** Immunofluorescence images of 12 months old WT, APP23 and A9KO (TLR9-deficient APP23 mice) stained for dsDNA (green) and nucleus (DAPI, blue) in the CA3 region of the hippocampus and in the entorhinal cortex. Scale bar: 10 μm. **(B)** Quantification of extranuclear dsDNA. n=6-7 images/mice analyzed for 12 months and n=4 images/mice analyzed for 18 months old mice (n=3 for all). ns: not significative, **p<0.01, ***p<0.001, ****p<0.0001; one-way ANOVA was performed with Tukey post-hoc test. **(C)** TNF-α production by microglia purified from adult WT and TLR9-KO mice, stimulated with 3 μg/ml of enriched mitochondrial DNA (mtDNA) from 15 months old WT or APP23 brain and measured by ELISA. n=3. ****p<0.0001; one-way ANOVA was performed with Tukey post-hoc test. mtDNA: mitochondrial DNA. **(D)** Immunofluorescence images of 18 months old APP23 mice stained for dsDNA (red, extranuclear: white arrows), microglia (IBA-1, green) and fibrillar β-sheeted Aβ plaques (thioflavin S) in the cortex. Magnification shows one microglia interacting with extranuclear dsDNA (yellow arrow). Scale bar: 50 μm. **(E)** Left: immunofluorescence images of 18 months old APP23 and A9KO mice stained for microglia (IBA-1, green), Aβ plaques (6E10, red) and nucleus (DAPI, blue) in the cortex. n=3 Scale bar: 10 μm. Right: IBA-1 relative fluorescence intensity (top) and microglia numbers (bottom) next to the plaques (<20 μm radius). A dot represents a pool of all plaques in one bregma level (n=4 levels) for IBA-1 fluorescence intensity and an individual animal for microglia number. ns: not significative; two-tailed student’s t-test was performed. **(F)** Left: immunofluorescence images of 18 months old APP23 and A9KO mice stained for PGRN (cyan) and Aβ plaques (6E10, magenta) in the cortex. n=3. Scale bar: 50 μm. Right: PGRN relative fluorescence intensity next to the plaques (<20 μm radius). One dot represents a pool of all plaques in one bregma level (n=4 levels). ***p<0.001; two-tailed student’s t-test was performed.

Microglia are the major immune and inflammatory cells of the brain and are recruited to Aβ plaque sites, which can in turn activate them and may aggravate AD pathology in the case of unsuccessful Aβ clearance. At 18 months of age, immunostaining with Iba1 antibody showed reduced Iba1 immuno-reactivity around Aβ deposits, detected with the 6E10 antibody, in brains of A9KO compared to APP23 mice but no change in the overall number of plaque-recruited microglia (Figure 1E). PGRN is a lysosomal resident or secreted glycoprotein encoded by the *Grn* gene that is cleaved to generate individual peptides and is associated with the modulation of inflammatory responses. Full-length secreted PGRN has anti-inflammatory and neurotrophic functions whereas GRN peptides were described to bind and facilitate the delivery of TLR9 ligand to the lysosome where TLR9 activation takes place (Park et al. 2011). In addition, mutations in *Grn* predispose to the development of AD (Lee et al. 2011) and increased PGRN expression is detected in the brain of AD patients and in mice developing AD, specifically in neurons and microglia (Minami et al. 2014, Mendsaikhan et al. 2019). To determine if PGRN is affected by TLR9 deficiency, we performed immunostaining of PGRN and showed significantly more PGRN clustering around Aβ plaques in the brain of APP23 mice relative to A9KO mice (Figure 1F).

All these results suggest that extranuclear dsDNA in the brain of APP23 is internalized by microglia and potentially triggers TLR9 activation.

### Phospholipase D3 co-localizes with dsDNA and TLR9 in microglia and is increased in the brain of APP23 mice

PLD3 is a lysosomal enzyme which degrades TLR9 ligands such as mtDNA to limit the accumulation of nucleic acids in the lysosome and is expressed by neurons and activated microglia in the CNS. As such, PLD3 regulates endosomal TLR9 sensing and *Pld3* gene polymorphism is linked to the development of AD. Because we found a strong accumulation of extranuclear dsDNA in APP23 mice, we assessed the expression of PLD3 in the brain of APP23 and A9KO mice. First, in both the hippocampus and entorhinal cortex, PLD3 strongly and preferentially co-localizes with TLR9 in TLR9-GFP mice (Figure S2A-B). We also saw rare occurrences of PLD3 and TLR9 colocalization in Iba1^+^ cells, as PLD3 is very weakly expressed in healthy microglia (Figure S2C). In pathological setting in these two regions, the proportion of cells expressing PLD3 was significatively increased in microglia and neurons from 12 months old APP23 mice in comparison to A9KO and WT mice (Figure 2A-D). Furthermore at 18 months, cells displayed increased PLD3 immuno-reactivity (Figure 2E-F) and dsDNA was localized in PLD3 positive compartments, especially in the entorhinal cortex of APP23 mice (Figure 2G-H). Bulk RNA-seq on purified CD11b^+^ cells also confirmed increased *Pld3* expression in APP23 microglia compared to WT at 18 months, an increase that was not statistically different between WT and A9KO cells (see later Figure S4D). A small increase in the proportion of non-microglial PGRN positive cells was also observed at 12 months of age (Figure 2A-B).

**Figure 2.**
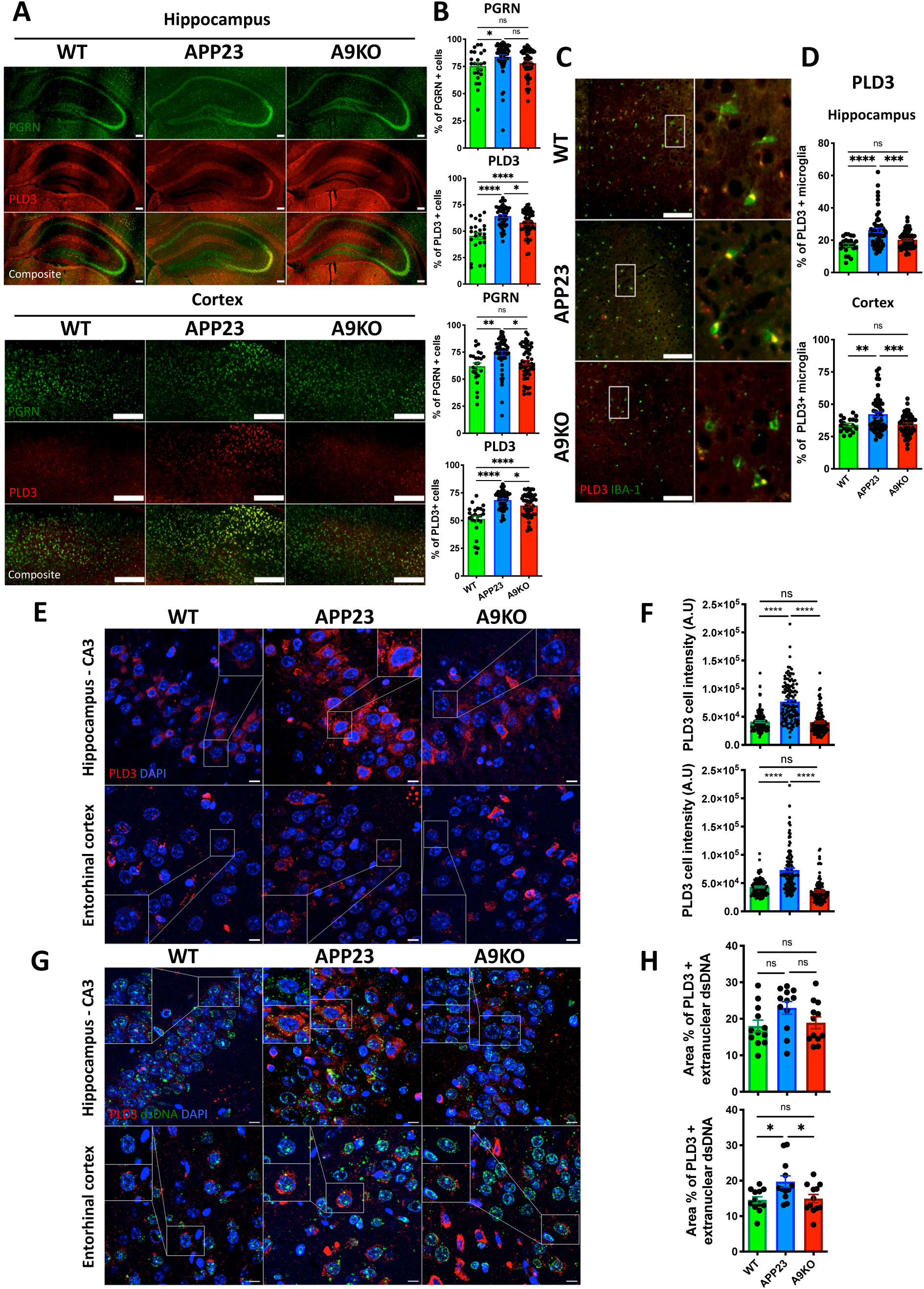
Expression of phospholipase 3 (PLD3) and progranulin (PGRN) are influenced by TLR9 during amyloid pathology. **(A)** Immunofluorescence images of the hippocampus and cortex from 12 months old WT (n=3), APP23 (n=6) and A9KO (n=6) mice stained for PGRN (green) and PLD3 (red). Scale bar: 100 μm. **(B)** Quantification of PGRN^+^ and PLD3^+^ non-microglial cells. Each dot represents one hippocampal or cortical region (n=8-10 regions per animal). ns: not significative, *p<0.05, **p<0.01, ****p<0.0001; one-way ANOVA was performed with Tukey post-hoc test. **(C)** Immunofluorescence images of the hippocampus from 12 months old WT (n=3), APP23 (n=6) and A9KO (n=6) mice stained for IBA-1 (green) and PLD3 (red). Scale bar: 100 μm. **(D)** Quantification of PLD3^+^ microglia. Each dot represents one hippocampal or cortical region (n=8-10 regions per animal). ns: not significative, **p<0.01, ***p<0.001, ****p<0.0001; one-way ANOVA was performed with Tukey post-hoc test. **(E)** Immunofluorescence images of 18 months old WT, APP23 and A9KO mice stained for PLD3 (red) and nucleus (DAPI, blue) in the CA3 region of the hippocampus and in the entorhinal cortex. n=3. Scale bar: 10 μm. **(F)** PLD3 relative cell fluorescence intensity. n=20 cells or more/animal/region were analyzed. n=3. ns: not significative, ****p<0.0001; one-way ANOVA was performed with Tukey post-hoc test. **(G)** Immunofluorescence images of 12 months old WT, APP23 and A9KO mice stained for PLD3 (red), nucleus (DAPI, blue) and dsDNA (green) in the CA3 region of the hippocampus and in the entorhinal cortex. n=3. Scale bar: 10 μm. **(H)** Quantification of PLD3^+^ extranuclear dsDNA. Each dot represents one image quantified (n=4 images per animal). ns: not significative, *p<0.05; one-way ANOVA was performed with Tukey post-hoc test.

Prosaposin (PSAP) interacts with PGRN and traffics with PGRN to the lysosome to support lysosomal functions (Paushter et al. 2018), notably by controlling cathepsin D activity (Valdez et al. 2017). PSAP is also involved in lysosomal lipid remodeling (Hiraiwa et al. 1992) and PGRN-PSAP interaction leads to PGRN sequestration at Aβ plaque sites, which may play a role in the early events of AD by diminishing the neuroprotective and anti-inflammatory functions of PGRN (Mendsaikhan et al. 2019). Overall expression of PSAP and PGRN was decreased especially in the hippocampus of 12 months old A9KO mice (Figure S2D-E). These results suggest that A9KO cells display decreased lysosomal stress that limits the requirement for PSAP-mediated lysosomal support and lipid remodeling.

Altogether, these findings indicate an increase in the expression of PLD3 in APP23 mice to degrade dsDNA which strongly accumulates over time which is necessary to limit TLR9 sustained activation.

### TLR9 deficiency restores cognitive defects and reduces Aβ plaque deposition in male APP23 mice

In order to determine if TLR9 deficiency affects Aβ pathology and cognitive deficits, we quantified Aβ plaque load by 6E10 immunostaining and assessed cognitive performance in memory-related tasks in 12 and 18 months old mice. We observed a significant decrease in the overall Aβ burden, the size and the number of individual plaques in male A9KO compared to male APP23 mice at 18 months of age, both in the hippocampus and in the cortex (Figure 3A-B). At 12 months, while Aβ load was very low for both genotypes, there was also a decreased number of plaques in A9KO male mice (Figure S3A-B). Surprisingly, female mice showed no significant difference in the extent of Aβ deposition at 18 months of age (Figure 3C-D) and A9KO mice had even more plaques at 12 months of age than APP23 mice (Figure S3A-B). Since the APP23 mouse model is also associated with neuronal loss specifically in the CA1 region of the hippocampus (Calhoun et al. 1998), we measured the pyramidal layer thickness and the number of nuclei in the CA1 region at 18 months of age. While APP23 mice displayed an increased CA1 neuronal loss compared to WT, this was not the case in A9KO mice (Figure S3C-D).

**Figure 3.**
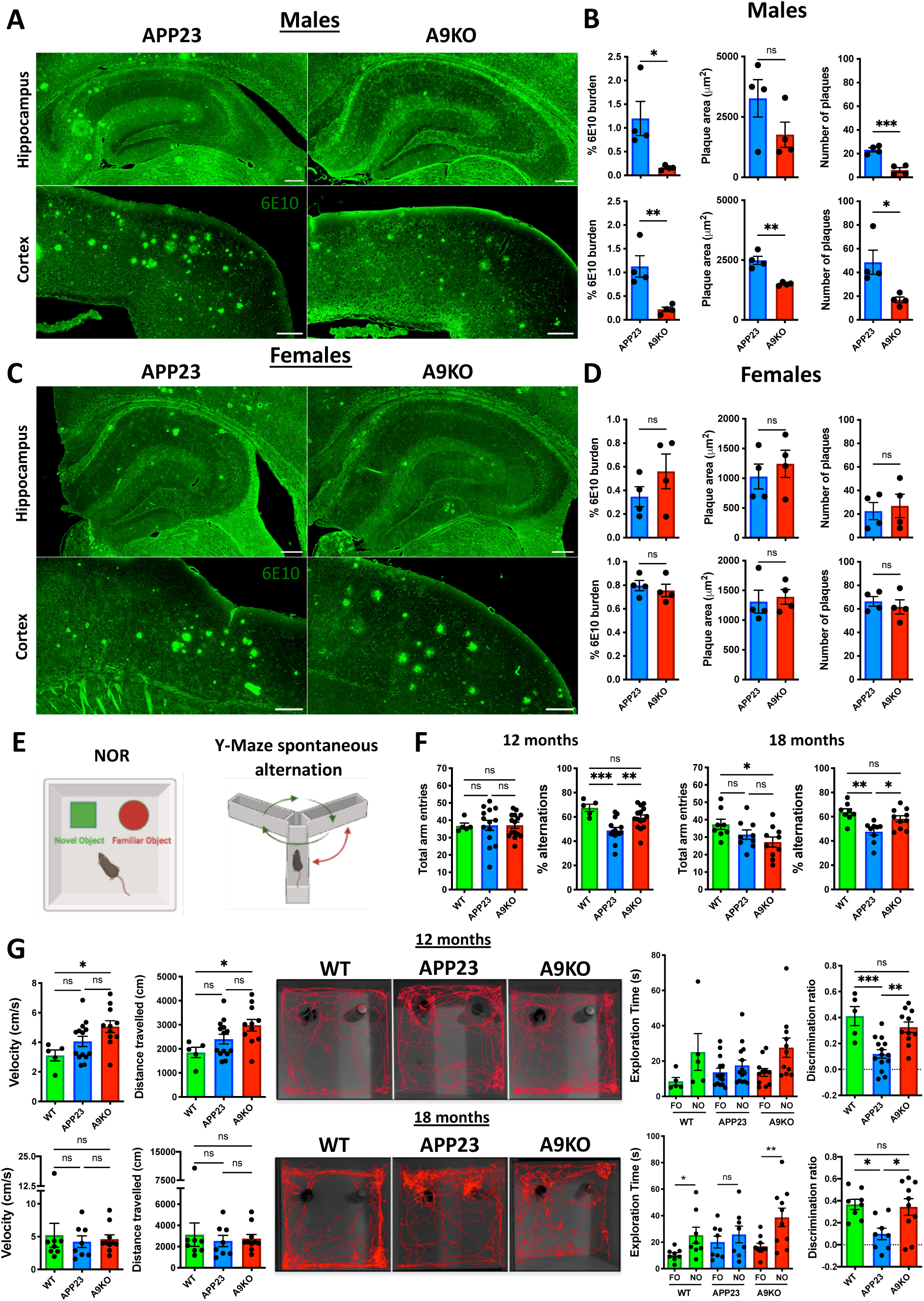
TLR9 contributes to cognitive deficits and amyloid pathology. **(A)** Aβ plaques stained with 6E10 antibody in the hippocampus and cortex regions of 18 months old male APP23 and A9KO mice. n=3. Scale bar: 200 μm. **(B)** Quantification of 6E10 burden, area of individual Aβ plaques and total number of plaques for male animals. One dot represents the pool of all plaques in one bregma level (n=4 levels). ns: not significative, *p<0.05, **p<0.01, ***p<0.001; two-tailed student’s t-test was performed. **(C)** Aβ plaques stained with 6E10 antibody in the hippocampus and cortex regions of 18 months old female APP23 and A9KO. n=3. Scale bar: 200 μm. **(D)** Quantification of 6E10 burden, area of individual Aβ plaques and total number of plaques for female animals. One dot represents the pool of all plaques in one bregma level (n=4 levels). ns: not significative; two-tailed student’s t-test was performed. **(E)** Schematic representation of the novel object recognition (NOR) test to assess long-term recognition memory and Y-Maze (spontaneous alternation) test to assess short-term spatial working memory. **(F)** Y-maze test. Left: number of total arm entries and % of alternations in 12 months old WT (n=5), APP23 (n=14), A9KO (n=15) mice. Right: same as in left in 18 months old WT (n=8), APP23 (n=9), A9KO (n=10) mice. Each dot represents an individual animal. ns: not significative, *p<0.05, **p<0.01, ***p<0.001; one-way ANOVA was performed with Tukey post-hoc test. **(G)** NOR test. Upper panel: velocity, distance travelled, representative animal tracks, object exploration time and discrimination ratio in 12 months old WT (n=5), APP23 (n=14), A9KO (n=11) mice. Lower panel: same as upper panel in 18 months old WT (n=8), APP23 (n=8), A9KO (n=10) mice. ns: not significative, *p<0.05, **p<0.01, ***p<0.001; one-way ANOVA was performed with Tukey post-hoc test; two-tailed student’s t-test was performed for the exploration time comparing NO to FO. NO: Novel Object, FO: Familiar Object.

Regarding behavior, at both timepoints analyzed, APP23 mice showed deficits in long-term recognition memory and short-term spatial working memory, as evidenced by the novel object recognition (NOR) and the Y-maze (spontaneous alternation) tests, respectively (Figure 3E-G). Mice lacking TLR9 expression performed significantly better than APP23 mice in both of these tests, with a stronger effect at 12 months of age (Figure 3F-G). While amyloid pathology was associated with anxiety-related behaviors in the light-dark box test, no difference was observed in locomotion or in depressive-like behaviors (Figure S3E). In order to exclude a direct effect of TLR9 deficiency on cognitive performance, we performed identical behavioral tests in healthy 9 months old WT and TLR9-KO mice, at a time point that includes both the cognitive deficits and the start of Aβ pathology seen in the APP23 mouse model. In the absence of amyloid pathology, TLR9 deficiency had no impact on memory-related or other cognitive functions (Figure S3F), suggesting that TLR9 involvement in these processes is dictated by progressive Aβ pathology.

Overall, we demonstrate that TLR9 participates to disease-associated cognitive deficits in both males and females and to Aβ deposition only in male mice.

### TLR9 deficiency drives microglial anti-inflammatory M2 reprogramming

Ageing, prolonged exposure of microglia to Aβ plaques and their associated products as well as local inflammatory cytokines lead to their activation and phenotypic transition from homeostatic to disease-associated (DAM) and type I interferon-responsive microglia (IRM) (Keren-Shaul et al. 2017, Sala Frigerio et al. 2019). To characterize immune cells involved in our model, we performed bulk RNA-seq on CD11b^+^ cells isolated from whole brain of 18 months old APP23, A9KO and WT mice. The majority of CD11b^+^ cells were microglia expressing specific markers such as *Tmem119* and *P2ry12* (Figure 4A). Of note, principal component analysis clustered microglia separately depending on the three genotypes, suggesting distinct transcriptional signatures (Figure S4A). Due to the overall variations in differentially expressed gene (DEG) alterations, we considered that a 40% up- or downregulation was relevant (with a Q-Value<0.05). Analysis of DEGs revealed more pronounced changes in microglia, with TLR9 deficiency leading to an overall transcriptional downregulation, in accordance with TLR9 potential to activate immune cells (Figure 4B and Figure S4B). Gene-set enrichment analysis showed that APP23 microglia had an enriched signature associated with myeloid and microglial cell ageing compared to A9KO (Figure 4C and Figure S4C). KEGG pathway analysis also revealed several enriched pathways in A9KO downregulated DEGs. These included oxidative phosphorylation and glycolysis suggesting an overall dampened metabolism, HIF-1 signaling pathway which has been linked to glycolytic metabolic shift and pro-inflammatory phenotype (Sun et al. 2024) as well as the mitophagy pathway (Figure 4D). In addition, pathways involved in glutamatergic synapse and alanine, aspartate and glutamate metabolism were also enriched in A9KO downregulated DEGs, suggesting a decreased involvement in excitatory synapse remodeling, a dysfunctional process occurring during AD. In CD11b^-^ cells, pathology induced an increase in the proportion of upregulated DEGs but TLR9 deficiency had no effect on this balance (Figure S4B) implying that TLR9 deficiency impacts mainly microglia. Analysis of microglial signature revealed four different phenotypes: homeostatic, anti-inflammatory M2a, DAM and IRM. Overlaying WT and A9KO genotypes showed that A9KO microglia maintained their level of several homeostatic genes such as *P2ry12, Tmem119, Tgfbr1, Txnip, Mef2a, Ccr5, Adgrg1* and *Gpr34*, in contrast to APP23 microglia which had an increase in the expression of genes related to DAM and IRM populations (Figure 4E). To validate increased expression of the DAM marker *Gpnmb* in APP23 mice, we co-immunolabelled the cortex for GPNMB, Iba1 and Thioflavin S and showed higher GPNMB expression in microglia associated to dense core Aβ plaques (Figure 4F). As IRM have been linked to the elimination of DNA-damaged neurons (Escoubas et al. 2024), the emergence of this profile also suggests an increased Aβ-associated toxicity to neurons in APP23 mice. Moreover, compared to APP23, A9KO microglia displayed increased expression of M2a phenotype markers such as *Chil3* (Ym1) and *Retnla* (Fizz1) among others, a subtype specialized in resolving inflammation and promoting tissue repair (Franco and Fernández-Suárez 2015) (Figure 4E). Moreover, while the M2a surface marker CD163 was not different between WT and A9KO, APP23 microglia displayed significant decreased expression compared to WT (Figure S4D). Interestingly, we also observed decreased *Sting1* expression in APP23 microglia suggesting that this pathway is not preferentially upregulated in microglia during disease progression in our model (Figure S4D). These changes in A9KO microglial phenotype were associated with an increase of anti-inflammatory IL-10 and FGF21 and anti-viral type III interferon cytokines in the serum at 12 months of age (Figure S5), indicating that TLR9 deficiency also impacts inflammation at the periphery.

**Figure 4.**
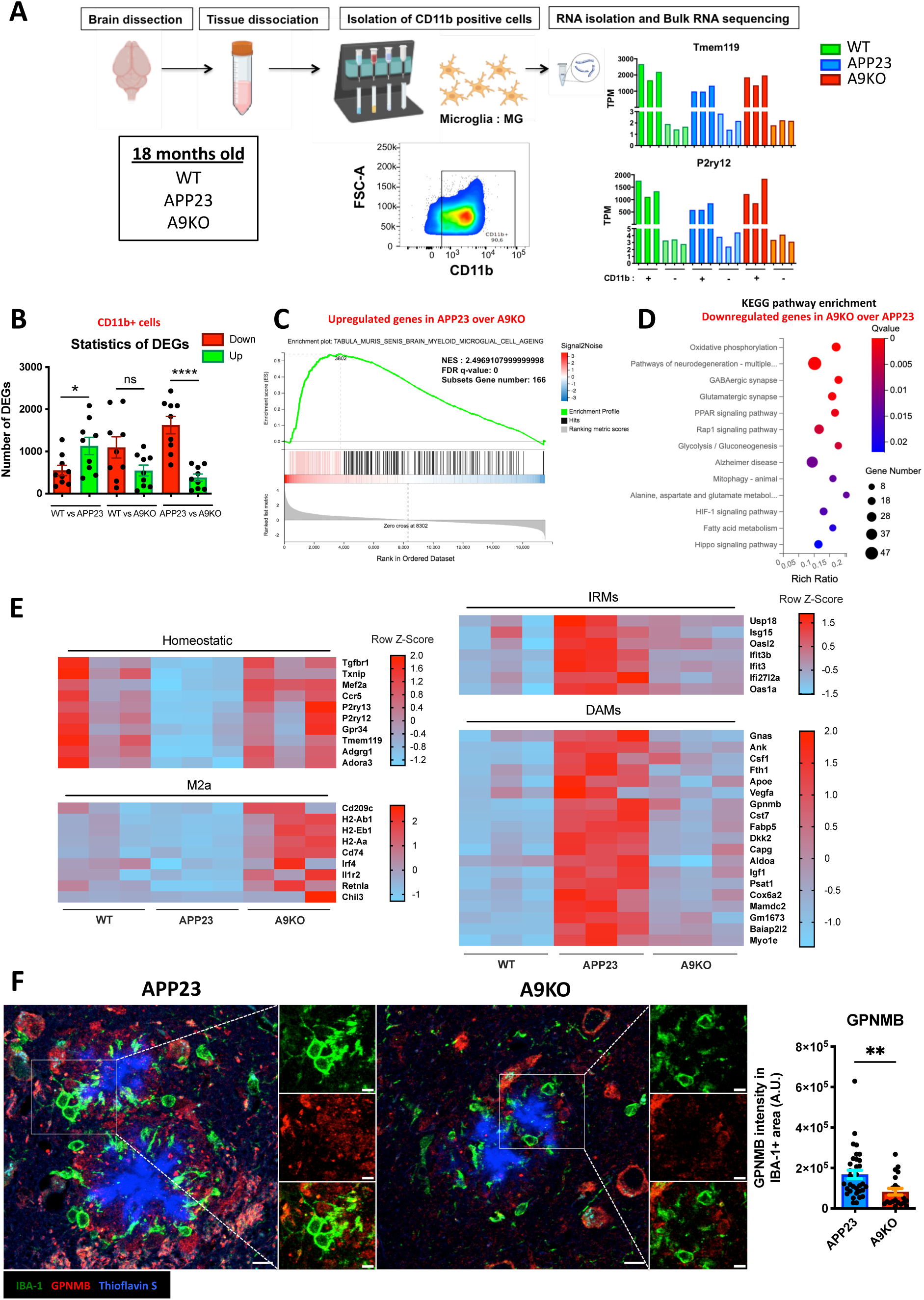
TLR9 expression controls microglial phenotype during AD. **(A)** Schematic representation of microglia isolation, purity (CD11b^+^) and enrichment (Tmem19 and P2ry12 high) to perform bulk RNA-seq in 18 months old WT, APP23 and A9KO mice. n=3. TPM: transcripts per million. **(B)** Numbers of up- and downregulated DEGs in CD11b^+^ samples. ns: not significative, *p<0.05, ****p<0.0001; two-tailed student’s t-test was performed. **(C)** Gene Set Enrichment Analysis (GSEA) performed on the upregulated genes in APP23 versus A9KO microglia. NES: normalized enrichment score, FDR: false discovery rate. **(D)** KEGG pathway enrichment analysis performed on downregulated genes in A9KO versus APP23 microglia. **(E)** Transcriptional analysis of homeostatic, M2a, type I IFN-responsive (IRM) and disease-associated (DAM) microglia genes in WT, APP23 and A9KO. Heatmap shows row Z-Score. **(F)** Left: immunofluorescence images of 18 months old APP23 and A9KO mice stained for Aβ plaques (thioflavin S, blue), microglia (IBA-1, green) and GPNMB (red) in the cortex. Scale bar: 10 μm, scale bar magnification: 5 μm. Right: GPNMB relative fluorescence intensity in plaque-associated microglia. 6 to 13 plaques per animal were analyzed for APP23 and A9KO. n=3. **p<0.01; two-tailed student’s t-test was performed.

Therefore, TLR9 deficiency attenuates microglial activation and induces microglia reprogramming to limit Aβ plaque toxicity and promote repair of Aβ-induced tissue damage.

### Lack of TLR9 expression restores lysosomal fitness

In AD, microglial switch from homeostatic to DAM phenotype may lead, if sustained, to lysosomal dysfunction, favouring defective clearance of Aβ and neurodegeneration. To address this, we analysed the expression of lysosomal-related genes in APP23, A9KO and WT microglia. Overall, lysosomal genes involved in phagocytosis, endosomal-lysosomal trafficking, macromolecule transport and regulation of lysosomal acidification were upregulated in A9KO compared to APP23 (Figure 5A). Among these, we noticed the two subunits of the signal regulatory protein β 1 (SIRPβ1) receptor, associated with Aβ uptake in microglia (Gaikwad et al. 2009). In contrast, genes participating in lysosomal quality control (LQC) and more specifically lysophagy (Koerver et al. 2019, Liu et al. 2020) such as *Fbxo2* and *Ube2ql1* were upregulated in APP23 compared to A9KO. *Bloc1s1*, a subunit of the biogenesis of lysosome-related organelle complex-1 (BLOC-1), associated with organelle biogenesis and function was also increased in APP23 microglia. Moreover, while the expression of two subunits of the V-ATPase, responsible for lysosomal acidification, were also upregulated in APP23 microglia, this change was accompanied by a significant but slight upregulation of *Ddrgk1* (∼30%), shown to mediate the degradation of V-ATPase subunits (Cao et al. 2021) in order to maintain a stable pool in the lysosome (Figure S4D). Interestingly, several transporters such as *Gpr55* (also known as LYCHOS), *Slc38a9* and *Slc4a7* were upregulated in A9KO microglia and were shown to control the activation of mTORC1 (Rebsamen et al. 2015, Ali et al. 2022, Shin et al. 2022), a pathway also involved in LQC and more specifically in the regulation of TFEB-mediated lysosome biogenesis. Because TFEB is a transcription factor that translocates to the nucleus to control lysosome biogenesis, we investigated whether TFEB activity was enhanced in APP23 microglia. Nuclear translocation of TFEB, necessary for its activity, was assessed by immunofluorescence in APP23 and A9KO microglia and no difference was detected at steady-state (Figure S6A-B). One of the genes upregulated in A9KO is cathepsin C (*Cstc*), a lysosomal dipeptidyl peptidase. Its activity in lysosomes is controlled by cystatin-F (Hamilton et al. 2008) (*Cts7*), a gene that is downregulated in A9KO microglia (Figure 4E). We assessed the recruitment of cathepsin C in lysosomes by immunofluorescence, where cathepsin C becomes fully functional and mature. The expression of cathepsin C in LAMP1^+^ lysosomes was significantly higher in A9KO than in APP23 microglia (Figure 5B-C). While secreted cathepsin C has been shown to promote microglia activation and neuroinflammation (Liang et al. 2016, Liu et al. 2019), these results suggest that this upregulation is related to its lysosomal functions. In line with alterations of lysosomal proteases, Cystatin B (*Cstb*), an inhibitor of lysosomal cathepsins B, L and S (Rinne et al. 2002) was also upregulated in APP23 microglia compared to WT, in contrast to A9KO microglia which showed no difference (Figure S4D).

**Figure 5.**
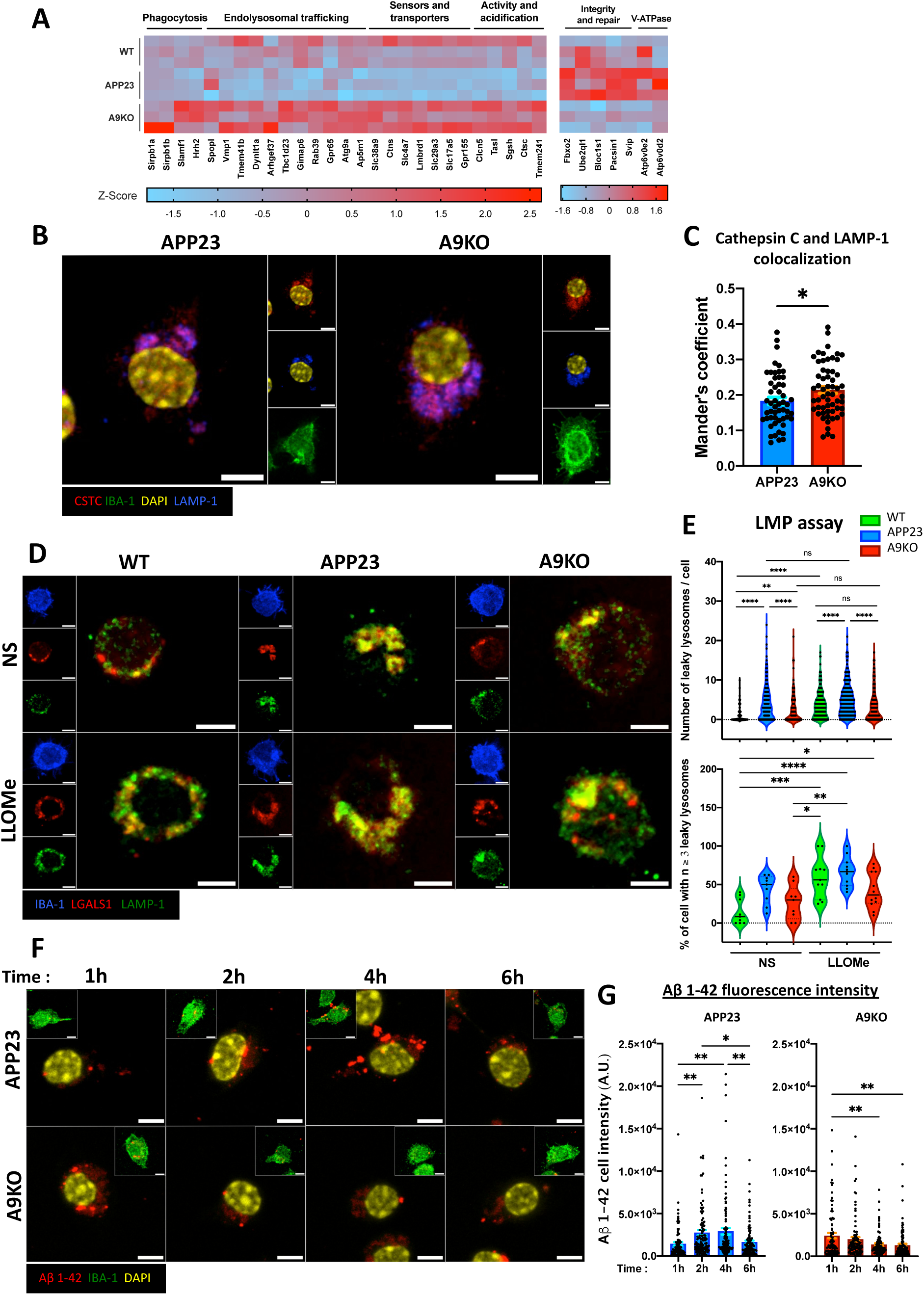
TLR9 promotes lysosomal integrity disruption in microglia during AD. **(A)** Transcriptional analysis of lysosomal-related genes in WT, APP23 and A9KO microglia. n=3. Heatmap shows Z-Score (for genes). **(B)** Immunofluorescence images of purified microglia from 10 to 15 months old APP23 and A9KO mice stained for IBA-1 (green), cathepsin C (red), lysosomes (LAMP-1, blue) and nucleus (DAPI, yellow). Scale bar: 5 μm. **(C)** CTSC and LAMP-1 colocalization expressed as Mander’s coefficient. n=20 cells or more per condition for each experiment were analyzed. n=2. *p<0.05; two-tailed student’s t-test was performed. CTSC: cathepsin C. **(D)** Immunofluorescence images of purified microglia from 10 to 15 months old WT, APP23 and A9KO mice stimulated or not with 0.5 mM LLOMe for 2 h and stained for microglia (IBA-1, blue), LGALS1 (red) and lysosomes (LAMP-1, green). Scale bar: 5 μm. LLOMe: L-Leucyl-L-Leucine Methyl Ester, LGALS1: galectin 1. NS: non-stimulated. **(E)** Quantification of the number of LGALS1^+^/LAMP-1^+^ puncta per cell (top) and of the percentage of cells with three or more puncta per field acquired (bottom). n= 20 cells or more per condition for each experiment was analyzed using IMARIS. n=3. NS: non-stimulated. ns: not significative, *p<0.05, **p<0.01, ***p<0.001, ****p<0.0001, one-way ANOVA was performed with Tukey post-hoc test. 4 Immunofluorescence images of purified microglia from 10 to 15 months old APP23 and A9KO mice incubated with 4 μM of Aβ 1-42 fibrils coupled to HilexaFluor 488 (red) for 1, 2, and 6 hours and stained for microglia (IBA-1 green) and nucleus (DAPI, yellow). Scale bar: 5 μm. **(F)** Aβ 1-42 relative intracellular fluorescence intensity in APP23 and A9KO microglia. n= 20 cells or more per condition for each experiment was analyzed. n=3. *p<0.05, **p<0.01; one-way ANOVA was performed with Tukey post-hoc test.

In microglia, the interplay between lysosomal dysfunction and inflammatory mechanisms can disrupt lysosomal integrity and exacerbate the aggregation of pathological proteins through defective microglial clearance. To dissect how the pathology affects lysosome integrity, we purified microglia from 10 to 15 months old APP23, A9KO and WT mice and performed lysosome permeabilization (LMP) assay and monitored microglial response kinetics to Aβ 1-42 peptides. The LMP assay consists in treating cells with L-leucyl-L-leucine methyl ester (LLOMe) which induces lysosomal membrane disruption without killing the cells (Aits et al. 2015). We treated microglia for 2 h and stained for galectin 1 (LGALS1) to assess its recruitment to leaky lysosomes. In the absence of LLOMe, microglia from APP23 mice showed more disrupted lysosomes compared to A9KO and WT as evidenced by the increased number of LAMP-1^+^ LGALS1^+^ puncta (Figure 5D-E). The treatment with LLOMe induced LMP, which was significantly more important in APP23 compared to A9KO and WT (Figure 5D-E), suggesting an increased susceptibility as well. Interestingly, plaque-associated PGRN was more present in recruited microglia in A9KO compared to APP23 mice (Figure S6C-D) suggesting that PGRN acts intracellularly to support lysosomal functions, as previously reported (Paushter et al. 2018). In line with decreased PGRN in plaque-associated APP23 microglia (Figure S6C-D), there was also a small but significant decrease (∼30%) in *Sort1* expression, coding for Sortilin, a protein that mediates the uptake of secreted PGRN (Hu et al. 2010, Liao et al. 2024) (Figure S4D). We then pulsed cells with fluorescently-labelled Aβ 1-42 peptides for different times and analyzed intracellular fluorescence intensity. At 2, 4 and 6h, APP23 microglia had increased intracellular Aβ 1-42 fluorescence compared to A9KO microglia. The peak of intracellular Aβ was reached at 1 h in A9KO microglia suggesting accelerated Aβ degradation (Figure 5F-G).

Overall, these findings indicate that APP23 microglia have impaired lysosomal fitness with increased damage to this organelle, which contributes to a defective Aβ clearance. Lack of TLR9 expression during disease progression restored lysosomal homeostasis.

### Extranuclear dsDNA accumulates in AD patients and modulation of TLR9 activation mediates the susceptibility to lysosomal disruption in human microglial cell line

We next investigated whether inhibiting or chronically inducing TLR9 activation with CpG-A alters lysosome fitness in human microglia. For this, we activated TLR9 in HMC3, a human microglia cell line, for 96 h, and performed the LMP assay described previously. The results showed that sustained TLR9 activation leads to increased susceptibility to LMP induction, with cells chronically-primed with CpG displaying an increased number of damaged lysosomes as demonstrated by the high number of lysosomal LGALS1^+^ puncta (Figure 6A-B, top panel). To inhibit TLR9, we used the lysosomotropic molecule AZP2006, which was shown to reduce inflammation *in vitro* and restore cognitive defects in a mouse model of AD by binding to the PSAP-PGRN complex and to inhibit human TLR9 signaling in HEK cells (Callizot et al. 2021). To confirm that AZP2006 inhibits TLR9 signaling also in microglia, HMC3 were treated with AZP2006 for 1 h, stimulated with specific TLR ligands and cytokine production was then measured by ELISA. As expected, AZP2006 inhibited intracellular TLR9 but not plasma membrane TLR4 (Figure 6C). We then asked whether this drug could impact lysosomal function. Incubation with AZP2006 significatively reduced lysosomal LGALS1^+^ puncta in cells treated with LLOMe alone and chronically stimulated with CpG for 96 h (Figure 6A-B, bottom panel). These results suggest that chronic activation or inhibition of TLR9 impacts the susceptibility to lysosomal permeabilization in human microglia.

**Figure 6.**
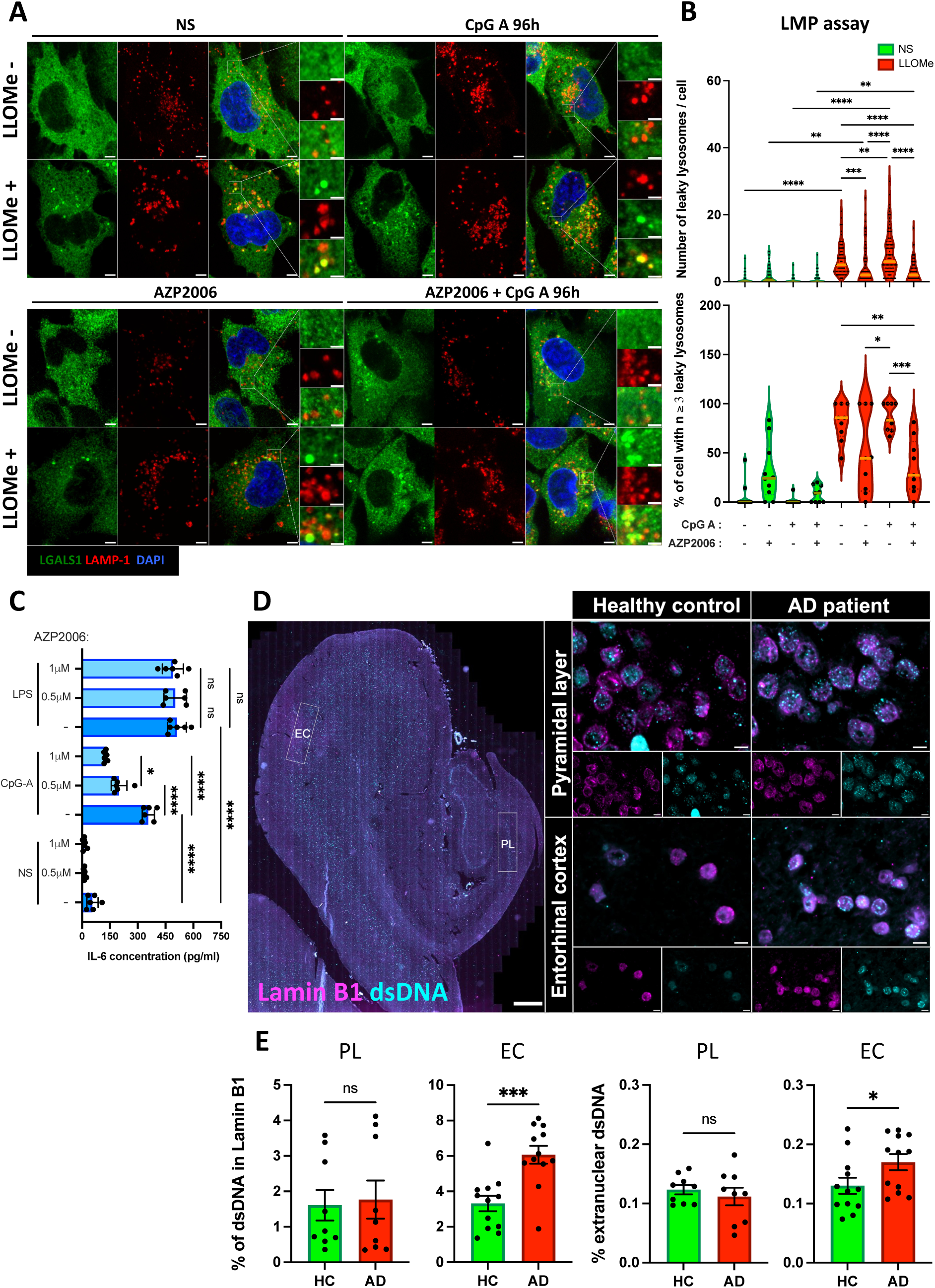
dsDNA accumulates in post-mortem brain from AD patients and TLR9 chronic activation or inhibition impacts lysosomal permeability in human microglia. **(A)** Immunofluorescence images of human microglia cell line (HMC3) cultured with CpG-A (2 μg/ml) for 96 hours, pre-incubated or not for 1 h with 0.5 μM of AZP2006, treated or not with 1 mM LLOMe for 2 h and stained for LGALS1 (green), lysosomes (LAMP-1, red) and nucleus (DAPI, blue). Scale bar: 5 μm. Scale bar magnification: 2 μm. **(B)** Quantification of the number of LGALS1^+^/LAMP-1^+^ puncta per cell (top) and of the percentage of cells with three or more puncta (bottom). n= 20 cells or more per condition for each experiment was analyzed. n=3. *p<0.05, **p<0.01, ***p<0.001, ****p<0.0001; one-way ANOVA was performed with Tukey post-hoc test. **(C)** IL-6 production by HMC3 cells pre-incubated or not for 1 h with 0.5 μM or 1 μM of AZP2006, stimulated or not with 1 μg/ml of CpG-A or 100 ng/ml of LPS and measured by ELISA. n=3. ns: not significative, *p<0.05, ****p<0.0001, one-way ANOVA was performed with Tukey post-hoc test. NS: non-stimulated. **(D)** Immunofluorescence images of the pyramidal layer of the hippocampus (top) and entorhinal cortex region (bottom) in human brain samples from a healthy control (n=3, left) and an AD patient (n=3, right) stained for dsDNA (cyan) and nuclear envelope (Lamin B1, magenta). On the left is displayed whole tissue with regions analyzed highlighted. Scale bar whole region: 1 mm. Scale bar insets: 5 μm. PL: pyramidal layer, EC: entorhinal cortex. **(E)** Quantification of the percentage of dsDNA area in the nucleus (within Lamin B1, left) or total percentage of extranuclear dsDNA for the field acquired (right) in the PL and in the EC. Each dot represents a quantified image (n=3 for PL and n=4 for EC for each individual). ns: not significative, *p<0.05, ***p<0.001; two-tailed student’s t-test was performed.

Finally, to verify the relevance of our findings in human concerning the release of dsDNA, we analyzed *postmortem* hippocampus and entorhinal cortex samples from AD patients and healthy controls. We immunostained sections with dsDNA and Lamin B1, the latter allowing to visualize the nuclear envelope. In accordance with our findings in mice, we observed an increased amount of extranuclear dsDNA in AD patients compared to healthy controls in the entorhinal cortex (Figure 6D-E). Altogether, these results show that AD patients display increased amounts of accessible TLR9 ligand which might lead to enhanced microglial activation.

## Discussion

Altogether, our data reveal a previously underappreciated role for TLR9 signaling in the course of AD progression. Whereas acute TLR9 signaling can prove beneficial to boost Aβ peptide clearance, we show that sustained activation of murine microglia through TLR9 signaling can contribute to cell activation and dysfunction, participating to the physiopathology of AD. For this, we used a slow-progressing model of amyloid pathology and focused on two timepoints, both the early and the late stage of Aβ deposition.

We show both in mouse and in human CNS that dsDNA, a potential ligand for TLR9 activation, accumulates extracellularly during AD, possibly due to age-associated genomic instability, Aβ-induced toxicity or defective mitophagy. We also demonstrate that murine microglia express TLR9 and respond to mtDNA in a TLR9-dependent manner, suggesting that increased dsDNA generation during AD may chronically activate microglia and promote neuroinflammation during AD progression. Moreover, only mtDNA isolated from APP23 elicited a response in WT microglia suggesting an exacerbated immunostimulatory potential induced by the pathology. More experiments are required to assess the characteristics of these ligands in pathological setting and how potential alterations can affect immune cell activation. Regarding AD physiopathology, we provide clear evidence that TLR9 contributes to AD progression as mice lacking TLR9 expression show less severe cognitive deficits and neuronal loss. In addition, since in males specifically, TLR9 deficiency led to a lower plaque load, we sought to determine changes in microglial phenotype and functions that could explain this difference in term of Aβ deposition control. As previously reported in AD mouse models (Keren-Shaul et al. 2017, Sala Frigerio et al. 2019), we observed a phenotypic switch in APP23 microglia that adopted the DAM and IRM phenotype, two microglia signatures associated with AD. In mice lacking TLR9, this shift was clearly attenuated suggesting that at least partially, TLR9 contributes to these phenotypic alterations. TLR9 deficiency did not only dampen this phenotypic switch but also promoted an M2a anti-inflammatory reprogramming, a subtype specialized in regulation of inflammation and in tissue repair (Franco and Fernández-Suárez 2025). As plaques induce cell toxicity and neuronal death, we suggest that DNA released from dying cells leads to chronic TLR9 activation *in situ*, thereby participating to microglial exhaustion, exacerbated inflammatory responses and defective Aβ degradation. Of note, this reprogramming was also associated with slight but significant changes in cytokine profile at the periphery in the early stages of the pathology, notably with increased IL-10, FGF21 and type III IFN cytokines in the serum. Studies reported that peripheral CpG administration promotes the mobilization of peripheral monocytes to control disease progression (Scholtzova et al. 2014). In contrast, we suggest that chronic activation of TLR9 in microglia leads to cell overactivation, impairing their ability to respond to and control Aβ deposition.

Other nucleic acid sensors participate to AD physiopathology through the modulation of myeloid cell activation and functions following mtDNA recognition (Shimada et al. 2012, Chung et al. 2024). For example, NLRP3 has been linked to microglial activation and lysosomal dysfunction, both participating to disease progression during amyloid deposition (Heneka et al. 2013, Friker et al. 2020). NLRP3 also contributes to defective Aβ clearance as its deletion or inhibition leads to a rewiring of microglial metabolism and promotes Aβ elimination (McManus et al. 2025). In addition, STING mediates the activation of microglia in the App^NL-G-F^/hTau-dKI mouse model and leads to exacerbated neuroinflammation and synaptic engulfment by microglia. In this model, Chung et al. showed that Aβ and tau-mediated toxicity triggers ectopic DNA accumulation in microglia that sustains STING activation. In addition, Thanos et al. showed that deleting STING reduces microglial activation, Aβ deposition and related cognitive deficits in the 5xFAD mouse model (Thanos et al. 2025). In our model, we did not observe a significant upregulation of NLRP3 and STING in APP23 microglia compared to WT. Even more surprising, STING was significantly downregulated in APP23 microglia compared to WT. While these data suggest that these pathways are not preferentially mobilized in microglia during slow-progressing amyloid pathology, more experiments are needed to confirm these results. As we also noticed that decreased STING expression is absent in A9KO microglia compared to WT, it is possible that STING signaling pathway may compensate for TLR9 deficiency. However, if such a compensatory mechanism is involved, it does not translate to adverse effects as microglial functions were still preserved and led to a better disease control in A9KO mice. Since STING and NLRP3 are cytosolic in contrast to lysosomal TLR9, subcellular localization of nucleic acid sensors as well as mechanisms that lead to dsDNA generation will be determinant to decipher the extent of immune responses involved and their impact on physiopathology. Moreover, the way immune cells internalize cell-free DNA and how endogenous or extracellular DNA is processed intracellularly will also determine which immune pathway is activated. Finally, as lysosome integrity is critical for both effective mitophagy and degradation of exogenous nucleic acids, disruption of lysosomal membrane would inevitably shift this balance in favor of cytosolic sensors such as STING. While we do not exclude that these receptors may play a role in microglia mediated-activation, more studies need to address the interplay of these different pathways during neurodegeneration and their integration to AD physiopathology.

Recent reports show that endosomal-lysosomal network (ELN) dysfunction contributes both to Aβ peptide generation by neurons and decreased clearance in microglia (Lee et al. 2022, Quick et al. 2023). TLR9 being a lysosomal protein, we thus investigated lysosomal changes in A9KO microglia. In parallel with the emergence of disease-associated microglial phenotypes, several genes involved in lysophagy were upregulated in the presence of TLR9. These included Ube2ql1 which is required for the ubiquitination of damaged lysosomes to perform lysophagy (Koerver et al. 2019). In addition, Fbxo2 was recently shown to be recruited to damaged lysosomes, acting as a member of the ubiquitin-ligase complex necessary to perform lysophagy, notably in the context of Niemann-Pick type C disease (Liu et al. 2020).

In contrast, TLR9-deficient microglia displayed increased expression of genes that support lysosomal homeostasis and functions, implying that sustained TLR9 signaling over time contributes to impaired lysosomal fitness. To address this, we purified microglia at late stage of the disease and performed LMP assay. Our results show that this transcriptomic signature translates to decreased LMP at steady-state and upon induction in TLR9-deficient microglia, in accordance with improved lysosomal integrity. In HMC3, chronic activation of TLR9 led to an opposing effect, increasing the susceptibility to LMP. Additionally, we used the lysosomotropic molecule AZP2006 that was previously reported to bind the PGRN-PSAP complex and stabilize PGRN levels. While the mechanisms involved are not clear, it was also linked to inhibition of TLR9 signaling, potentially as a bystander effect of PGRN stabilization (Callizot et al. 2021). We first confirmed that AZP2006 treatment leads to TLR9 signaling inhibition by assessing TLR9-mediated cytokine production in HMC3 cells. Because PGRN is crucial for lysosomal fitness and function, we investigated whether AZP2006 treatment could translate to decreased susceptibility for LMP upon induction. Our results show that AZP2006 effectively protects from LMP both in the absence of additional trigger and in the case of sustained TLR9 activation, thus counteracting TLR9-mediated increased LMP susceptibility. As Aβ clearance is dependent on lysosomal fitness, we also monitored Aβ 1-42 peptides expression following internalization in microglia. We show that this increased lysosomal fitness in TLR9-deficient microglia translates to more rapid degradation of Aβ 1-42 peptides.

PLD3 and PGRN are known risk factors for AD and their contributions to AD physiopathology is still unclear (Nackenoff et al. 2021, Rhinn et al. 2022). As PLD3 and PGRN both regulate TLR9 activation, our project provided a unique setting to study the impact of TLR9 expression on both of these proteins during Aβ deposition. First, to further support the idea that TLR9 signaling pathway contributes to cell activation, we demonstrate that dsDNA is localized in PLD3 positive compartments at late stage of the pathology, in line with its exonuclease function required to reduce TLR9 ligand availability and subsequent activation. Mice expressing TLR9 also displayed an increased expression of PLD3 in CNS-resident cells, including microglia. Moreover, RNA-seq analysis revealed that PLD3 is significantly upregulated in APP23 microglia compared to WT, which was not the case for A9KO mice. As PLD3 has been linked to the Activated Response Microglia (ARM) phenotype that also emerges during AD (Sala Frigerio et al. 2019), we propose that TLR9 activation can partially contribute to this phenotypic switch. In addition, we investigated PGRN dynamics in the CNS and plaque microenvironment as it was shown that PGRN expression in microglia supports lysosomal functions, notably through the regulation of bis(monoacylclycero)phosphate (BMP), a critical lysosomal phospholipid that promotes lysosomal hydrolases activity such as glucocerebrosidase (Logan et al. 2021), and attenuates plaque deposition. As PGRN products are also able to potentiate TLR9 activation, we also addressed whether PGRN expression and dynamics are affected by TLR9 deficiency. While PGRN overall expression in the CNS was not dramatically affected, plaque-associated PGRN was elevated in TLR9-expressing mice. There was also a shift in PGRN dynamics in the absence of TLR9, with plaque-associated PGRN being more present in microglia, in line with increased lysosomal fitness observed in A9KO microglia. In accordance with decreased microglial PGRN expression at the plaque site, APP23 microglia also displayed lower Sortilin expression which mediates the uptake of extracellular PGRN. While more studies are needed to specify PGRN and PLD3 involvement in AD progression, we provide strong arguments that these functions may be considered together with TLR9 signaling pathway.

Overall, our findings identify the nucleic-acid sensor TLR9 as a relevant immune regulator in the CNS. As TLR9 sustained activation leads to exacerbated immune responses and cellular dysfunction, we suggest that TLR9 is a pertinent target in the treatment of AD. Mitigating TLR9 signaling in microglia may offer an opportunity to dampen microglial dysfunction and promote disease control, bringing hope to the development of immune-based alternative therapies for AD patients.

## Limitations of the study

We found that TLR9 sustained signaling has detrimental effects on microglial activation states, lysosomal homeostasis and Aβ responses which was associated with less severe plaque load in males specifically. This discrepancy between male and female concerning Aβ burden implies that differential mechanisms may be involved. Moreover, as TLR9 signaling pathway was recently shown to be active in neurons and participate to cognition and learning processes (Jovasevic et al. 2024), it is possible that in our mouse model of progressive intracellular APP accumulation, neuronal TLR9 signaling may be detrimental in regard to APP metabolism. Since TLR9 was also shown to sustain autophagic flux (Lim et al. 2016), exacerbated TLR9 signaling in this setting may worsen ELN bottle-necks and participate to Aβ peptide generation. Moreover, as male and female microglia are different in terms of activation, inflammatory responses and phagocytic capacity (Lopez-Lee et al. 2024), the impact of TLR9 on microglia activation during AD progression may also be gender-specific. Larger cohorts with mice of both sexes will be required to answer these questions. Moreover, as we are in a model of slow amyloid deposition, mechanisms contributing to Aβ accumulation may differ in faster progressing models and we are now addressing these questions.

Additionally, more experiments are required to assess the degree of neuroinflammation and the quantitative changes of specific mediators that are present in the CNS as well as their correlation to disease progression. Given that TLR9 activation can lead to type I IFN production, a mediator for which IRMs are sensitive and that type I IFN is particularly important in the CNS and associated with AD (Lopez-Atalaya and Bhojwani-Cabrera 2025), determining the extent of type I IFN signaling in the CNS during amyloid deposition requires further attention. Moreover, at late stage of the disease, we established that TLR9-deficient microglia have adopted an anti-inflammatory phenotype. Whether this phenotype has emerged earlier and played an active part in limiting exacerbated inflammatory responses remains to be elucidated. This phenotype might as well be the consequence of improved ability to control plaque deposition and be needed posteriorly to promote the resolution of inflammation and tissue repair once microglia have cleared Aβ species. A more longitudinal approach would allow us to decipher these mechanisms.

It is known that dsDNA can activate TLR9 signaling and we showed that extranuclear dsDNA generation is a feature of AD. Whether this is related to defective mitophagy (Pradeepkiran and Reddy 2020), exacerbated cell death due to Aβ toxicity or increased neuronal DSBs and leakage through compromised nuclear envelope remains to be determined. In some instances, such as neuronal long-term potentiation, it was shown that released dsDNA due to DSBs displays increased frequency of GC-containing sequences, thereby providing adequate ligands for TLR9 signaling (Jovasevic et al. 2024). Moreover, while we show that this potential ligand for TLR9 is found in the plaque vicinity and that only mtDNA from APP23 mice can induce TLR9-dependent cytokine secretion, we think that more experiments are required to test its nature and how potential alterations may modulate microglia activation. Finally, while AZP2006 inhibits TLR9 signaling, protects from LMP and mirrors our setting of TLR9 deficiency, its mechanisms of action remain to be fully elucidated.

Regarding cell-specific TLR9 involvement, we show strong transcriptomic and functional alterations of *ex vivo* enriched microglia, suggesting a prominent role for these immune cells in AD progression. However, purified CD11b^+^ cells may contain a small fraction of infiltrating monocytes, which might participate in the phenotype we observe. To address this, we are generating a mouse model lacking TLR9 specifically in microglia.

## Material and methods

### Mice and human samples

For *in vivo* animal studies, all mice were on C57BL/6J background. Hemizygous APP23 mice carrying the Swedish double mutation under the control of the neuron-specific mouse Thy-1 promoter (APP_751_ *K670N/M671L) were obtained from Jackson laboratory (030504). APP23 mice were crossed to TLR9-KO (S. Akira) to generate APP23xTLR9-KO (referred to as A9KO mice). Non-APP23 transgenic littermates were used as control during the studies. All mice were genotyped for TLR9 deletion and APP23 transgene and a second genotyping was performed at the time of sacrifice. Mice were housed in groups according to the standard conditions at Charles Foix animal facility (Paris) under specific pathogen free (SPF) conditions, at constant room temperature (22°C) with a controlled 12 h day-night cycle and had access to food and water *ad libidum*. For behavioral analysis, mice with mixed genders were used at 12 and 18 months of age. Healthy TLR9-KO animals and WT littermates were used at 9 months of age. For functional assays, mice were sacrificed between 10 and 15 months of age. In addition, TLR9-GFP (M. Brinkmann) were housed at Necker Institute according to the standards conditions at the Specific Opportunist Pathogen Free (SOPF) institute animal facility (22°C, 12 h day-night cycle and access to food and water *ad libidum*). These mice were used only for free-floating immunohistofluorescence (IHF) experiments. Animal care and handling were performed in accordance with ethical guidelines from the European Convention for the Protection of Vertebrates Animals and all studies were performed in agreement with experimental protocol approval by the French ministry of Research (APAFIS n°136).

For human studies, paraffin-embedded sections from *postmortem* brain samples of histologically confirmed cases of AD (hippocampal region: CA1 to CA3 and entorhinal cortex) were obtained from brains collected in a Brain Donation Program of the Brain Bank “NeuroCEB” run by a consortium of Patients Association that include Fondation Vaincre Alzheimer, France Parkinson and ARSEP (Association for Research on Multiple Sclerosis). The signed consents were either from the patients or their next of kin in their name, in line with the French Bioethical laws. The three patients were at Braak stage VI with a Thal amyloid phase of 5, displayed type 2 CAA with ages of death of 62, 65 and 69. They were compared to three healthy sex-matched controls with an age of death of 31, 51 and 94. All samples were used for immuno-staining and acquisition on a slide scanner.

### Microglia cell culture

Microglia were purified from adult mouse brain using the Adult brain Dissociation kit (Miltenyi) and CD11b Microbeads (Miltenyi), following the manufacturer’s instructions. Briefly, mice were sacrificed by cervical dislocation, whole brain was isolated and put in ice-cold PBS. Meninges and olfactory bulbs were removed and all subsequent steps on the cortices were carried out at 4°C. After dissociation, debris removal and red blood cell lysis steps, cell suspensions were incubated with CD11b Microbeads and CD11b^+^ cells were then purified as previously described (Milner et al. 2022) using LS columns and QuadroMACS Separator (Miltenyi). After purification, CD11b^+^ suspensions were centrifugated at 300g for 10 min and maintained in culture for one day before experimental procedures. For functional experiments, *ex vivo* enriched microglial cells were cultured in DMEM (Dutscher) supplemented with 10% FCS, 1% penicillin/streptomycin (Dutscher) 1% L-glutamine (Dutscher) with the addition of 50 ng/ml GM-CSF (Thermofisher PeproTech) and 100 ng/ml M-CSF (Thermofisher PeproTech) (cDMEM) at 37^°^C in humidified 5% CO2, 95% air. The human microglial cell line HMC3 (isolated from the brain of a patient and established through SV40-dependent immortalization, ATCC #CRL-3304^TM^) was cultured in DMEM supplemented with 10% FCS, 1% penicillin/streptomycin, 1% L-glutamine and used between passage P5-P10. Cells were maintained at 37^°^C in humidified 5% CO2, 95% air and media was changed every 3 days until cells were used.

### Flow cytometry

Cells were washed in FACS buffer (PBS supplemented with 4% FCS), incubated for 15 min with Fc Block (BD Biosciences) and stained for 1 h on ice with Zombie red (Biolegend) and CD11b APC-eFluor 780. Stained cells were acquired with a BD LSR Fortessa flow cytometer and analyzed using FlowJo (Treestar) software. Plots show CD11b^+^ cells gated on live cells (Zombie Red^-^).

### Isolation of mtDNA and qPCR to assess enrichment

Two 15 months old WT or APP23 mouse brain were processed using the Adult brain Dissociation kit (Miltenyi). Cells were pooled by genotype and mitochondria were isolated using the Mitochondria Isolation Kit for Mammalian Cells (ThermoFisher Scientific). DNA was extracted using KAPA express extract (Sigma-Aldrich) and DNA concentration was measured using Nanodrop spectrophotometer (ThermoFisher Scientific). For control, genomic DNA (tail) from WT or APP23 mice was isolated using PureLink Genomic DNA kit (Invitrogen). Quantitative PCR was performed on both tail DNA and enriched mitochondrial DNA extracts diluted over 1000-fold range using D-loop primers at 0.3 μM concentration and β-actin primers at 0.1 μM concentration. qPCR reactions were run with the Power SYBR Green PCR Master Mix (ThermoFisher Scientific) on a C1000 Touch Thermal Cycler (Bio-Rad) with the following amplification profile: one cycle (95°C 10 min) and then 50 cycles (95°C 15 sec, 64°C 15 sec, 68°C 15 sec) followed by the melting cycle. Abundance of mitochondrial D-loop was normalized to the nuclear gene *Actb* using the 2–ΔΔCt method. Below are the primer sequences used.

D-loop: F. 5’-AATCTACCATCCTCCGTGAAACC- 3’
  R. 5’ -TCAGTTTAGCTACCCCCAAGTTTAA- 3’
β-actin: F. 5’ -GCTGTGCTGTCCCTGTATGCCTCT- 3’
  R. 5’ -CTTCTCAGCTGTGGTGGTGAAGC- 3’

### Cell treatments and LMP assay

For assessment of cytokine production, murine microglia or HMC3 cells were plated in 96 well plate (7.10^4^ cells for murine microglia and 5.10^5^ cells for HMC3) and left to adhere overnight at 37°C in the CO2 incubator. Cells were treated with 1 μg/ml class B CpG ODN 1826 (Invivogen), 3 μg/ml of enriched mitochondrial DNA, 1 μg/ml of class A CpG ODN 2216 (Invivogen) or with 100 ng/ml of LPS-EB Ultrapure (Invivogen) for 6 h at 37°C in the CO2 incubator. Cytokines were measured using ELISA kits (Thermofisher) and absorbances were determined with a Microplate Mithras LB 940 plate reader (Berthold). For the LMP assay, purified microglia were plated on Poly-D-lysine (Sigma-Aldrich) coated Ibidi μ-Slide VI 0.4 (Ibidi) and left to adhere overnight. HMC3 were also plated on Ibidi μ-Slide VI 0.4, stimulated or not with 2 μg/ml of class A CpG ODN 2216 (Invivogen) for 96 h at 37°C (every 24 h). Murine microglia and HMC3 were then incubated with 0.5 mM or 1 mM LLOMe (Santa Cruz) respectively for 2 h and then washed three times with PBS. For HMC3, AZP2006 (provided by Alzprotect) was pre-incubated at a concentration of 0.5 μM (0.5 μM and 1 μM for ELISA) for 1 h prior to LLOMe treatment. For conjugated Aβ treatment, β-Amyloid (1-42), Hilexa Fluor^TM^ 488-labeled peptides (Anaspec) were reconstituted as indicated by the manufacturer to a stock concentration of 20 μM and kept at -20°C until use. 24 h before treatment, peptides were diluted in PBS at a concentration of 4 μM and incubated at 37°C and in the dark to obtain fibrils. Cells were treated or not for 1, 2, 4 and 6 h at 37°C in the dark and in the CO2 incubator. Cells were then washed three times with PBS.

### Immunofluorescence

Following treatment, cells were fixed with 4% paraformaldehyde (PFA) (Thermofisher) for 10 min at room temperature (RT), washed once with PBS and then quenched with 0.2 M Glycine (Sigma-Aldrich) for 3 min. Cells were incubated with permeabilization/blocking buffer (1X PBS, 2% BSA (Sigma-Aldrich), 0.25% Triton X-100 (Sigma-Aldrich)) for 30 min at RT. Cells were then incubated overnight at 4°C with a combination of the following primary antibodies diluted in blocking buffer (BB) (1X PBS, 2% BSA): anti-galectin 1 antibody (EPR3205) (Abcam, #ab108389), anti-mouse CD107a (LAMP-1) clone 1D4B (Invitrogen, #14-1071-85), anti-human CD107a (BD Pharmingen, #555789), anti-AIF/Iba1 Antibody (Biotechne #NB100-1028), anti-Iba1 antibody (for immunocytochemistry) (Fujifilm Wako chemicals #019-19741), anti-TFEB antibody (Proteintech, #13372-1-AP) and anti-cathepsin C/DPPI antibody (Biotechne, #AF1034). Cells were then washed twice with BB and incubated for 30 min at RT with a combination of corresponding secondary antibodies diluted in BB: donkey anti-goat Alexa Fluor™ 488 (ThermoFisher, #A-11055), donkey anti-mouse Alexa Fluor™ 488 (ThermoFisher, #A-21202), donkey anti-rabbit Alexa Fluor™ 555 (ThermoFisher, #A-31572), donkey anti-mouse Alexa Fluor™ 594 (ThermoFisher, #A-21203), donkey anti-rabbit Alexa Fluor™ 594 (ThermoFisher, #A-21207), chicken anti-rat Alexa Fluor™ 647 (ThermoFisher, #A-21472) and donkey anti-rabbit Alexa Fluor™ Plus 647 (ThermoFisher, #A32795TR). After washing the cells twice with PBS, cells were incubated with DAPI (Abcam, 1 μg/ml in PBS) for 5 min at RT. Cells were then fixed again with 4% PFA for 5 min at RT. Finally, cells were incubated with 50 mM NH4Cl (Sigma-Aldrich) for 5 min at RT. Ibidi mounting medium (Ibidi) was applied and slides were either imaged right away or left at -20°C until acquisition the following day. For all experiments, all buffers were prepared fresh and filtered with 0.2 μm filters prior to use.

### Serum cytokines

Prior to perfusion, blood was collected from 12 months old animals by cardiac puncture in the left ventricle and left to clot at RT for approximatively 20 min. Tubes were then centrifugated at 2000g for 10 min at 4°C and serum was separated from the blood clot and collected. Samples were left at -20°C until processing. All samples were sent to Olink for quantification of cytokine levels (panel Target 48 mouse cytokines).

### Tissue preparation

Mice were anesthetized intraperitoneally with a mixture of ketamine/xylazine and transcardially perfused with ice-cold PBS and then ice-cold 4% PFA (no PFA perfusion for brains used for RNA-seq). Whole brain was isolated and for 18 months old animals, the two hemispheres were dissected in order to perform both bulk RNA-seq and quantification of Aβ pathology. For IHF, whole brain or brain hemispheres were put in 4% PFA at 4°C for 24 h. Tissue was then placed in a 30% sucrose solution containing 0.02% sodium azide (Sigma-Aldrich) until tissue sunk and then embedded or not in paraffin for tissue sectioning. Sections were either cut on vibratome (Leica) (40μm thickness) for free-floating IHF (TLR9-GFP mice) or on Microtome (Leica) (8μm thickness) for paraffin-embedded tissue (WT, APP23 and A9KO).

### Bulk RNA-seq

RNA was isolated from CD11b^+^ or CD11b^-^ cells using RNAspin Mini RNA isolation kit (Cytiva) following the manufacturer’s instructions. RNA samples (minimal input of 200 ng/sample) were sent to BGI Tech solutions for bulk RNA-seq. RNA library was prepared using DNBSEQ sequencing platform with PE100 as sequencing read length and 40 million reads per sample. For CD11b^+^ cells, a ribodepletion was performed prior to RNA library preparation with the same parameters. After sequencing, raw data was filtered with SOAPnuke software and all samples reached a minimum Q30 read quality of 94%. Clean reads were aligned to the mouse genome (GCRm39) using HISAT (Hierarchical Indexing for Spliced Alignment of Transcripts) software. DEGs were detected using the DESeq2 algorithm and analyzed on Dr.Tom software. For differential expression, DEGs with a log2FC of 0,486 and - 0,74 (40% upregulation or 40% downregulation, respectively) were considered as meaningful and the Q-value cut-off was set at 0.05. Only DEGs that passed these cut-offs between genotypes were shown in the figures. For volcano plots, all genes with a Q-value lower than 0.05 (-log10 (Q-value)>1.3) were shown. Similarly, for KEGG enrichment analysis, only pathways with a Q-value lower than 0.05 were shown. Gene-Set Enrichment Analysis was performed using the m8 module (cell type signature gene sets curated from cluster markers identified in single-cell sequencing studies of mouse tissue) from the Molecular Signatures Database (MSigDB).

### Image acquisition and colocalization analysis

Images were acquired on a confocal Leica SP8 gSTED microscope. All images were acquired as 3D Z-stacks either with 63x magnification (objective 63x HC PL APO OIL, NA=1.40) for immunofluorescence (IF) or with 40x magnification (objective 40x HC APO OIL, NA=1.30) for IHF. For whole section acquisition, tissue was acquired as a 7 μm (mouse tissue) or 5 μm (human tissue) stack on Olympus SLIDEVIEW VS200 scanner (Olympus corporation) with a 40X magnification (NA=0.95, 0.163 μm/pixel). For each experiment, all images shown were processed in the same manner and identical brightness/contrast parameters were set between conditions. To assess colocalization, settings on the microscope were set in order to have a 70 nm pixel resolution and single channel images were segmented using Ilastik software. On Fiji, regions of interest (ROI) for each cell were drawn and colocalization was expressed as Mander’s coefficient using JaCoP plugin. This process was automated using macros designed by Nicolas Goudin (INEM, Paris).

### LMP assay analysis using IMARIS

For LMP assay analysis, raw 3D Z-stacks images were subjected to the Pure Denoize plugin (4 cycles, 3 frames) on Fiji in order to reduce background before analysis on Imaris software. On Imaris, spot detector was used on LGALS1 and LAMP-1 denoized channels in order to detect LGALS1 or LAMP-1 puncta respectively. Object-Object statistics was selected in order to analyze the distance between galectin 1 and LAMP-1 puncta. Briefly, depending on overall size of puncta, estimated XY diameter and PSM elongation values were set for each image in order to detect all respective puncta. After detecting all puncta (adjustment of “Quality” parameter), the “absolute intensity” detection method was used in order to adjust the size of created objects in accordance with signal. For HMC3 cells, in the filter section, created objects were filtered first by volume (0.3 μm^3^) for both LAMP-1 and galectin puncta and were then subsequently filtered using “shortest distance between objects” with “below 300 nm” as a threshold. Volume-filtered galectin puncta was considered as the object with the shortest distance to volume-filtered LAMP-1 puncta for the analysis. For purified microglia, because of variability between experiments, threshold used ranged from 0.3 to 0.6 μm^3^ for the volume and from 300 to 500 nm for the “shortest distance between objects”. The same parameters were consistently used to compare different conditions within the same experiment. After these two object filtering steps, puncta were attributed to each individual cell in the field acquired using IBA-1 channel to delineate cell cytoplasm. Cells that were not entirely included in the acquired field were excluded from the analysis. The percentage of cells with 3 or more puncta in the acquired field was calculated.

### IHF on paraffin-embedded mouse and human tissue

Whole brain sections or single hemisphere sections from 12 or 18 month old mice were processed. Sections were first deparaffinized with 5 min incubation (two times) in Xylene (Sigma-Aldrich) and rehydrated by successive incubation in ethanol (EtOH) baths with decreasing concentrations of ethanol (Xylene+EtOH 100%, EtOH 100% x2, 75%, 50%, 25%, 15%, distilled water). For Aβ plaque antigen retrieval, sections were incubated for 10 min at RT in 98% formic acid (Sigma-Aldrich) and then rinsed two times for 1 min in distilled water. Standard antigen retrieval was then performed with citrate buffer (10 mM citrate, 0.05% Tween 20, pH 6.0) for 10 min at 95°C. After 30 min of cool-down, slides were washed for 5 min in PBS and permeabilized (PBS, 0.25% Triton X-100) for 10 min at RT, then washed two times for 5 min (TBS, 0.025% Triton X-100) and finally blocked in (TBS, 1% BSA, 0,3 M Glycine) for 2 h at RT. Sections were then incubated with the following primary antibodies overnight at 4°C: purified (azide-free) anti-β-Amyloid, 1-16, clone 6E10 (Biolegend, #803001), anti-β-Amyloid, 17-24 clone 4G8 (Biolegend, #800708), anti-DNA (Merck, #CBL-186), anti-AIF/Iba1 antibody (Biotechne #NB100-1028), anti-PGRN antibody (Bio-techne, #AF2557), anti-PSAP antibody (Proteintech, #10801-1-AP), anti-PLD3 antibody (Proteintech, #17327-1-AP), anti-MAP2 antibody (Proteintech, #17490-1-AP), anti-GPNMB antibody (Proteintech, #66926-1-Ig) and anti-TOM20 antibody (Proteintech, #11802-1-AP). After three 5 min washes, sections were incubated in the dark with the following secondary antibodies for 1 h at RT: donkey anti-goat Alexa Fluor™ 488 (ThermoFisher, #A-11055), chicken anti-rabbit Alexa Fluor™ 488 (ThermoFisher, #A-21441), donkey anti-mouse Alexa Fluor™ 488 (ThermoFisher, #A-21202), donkey anti-rabbit Alexa Fluor™ 555 (ThermoFisher, #A-31572), donkey anti-mouse Alexa Fluor™ 555 (ThermoFisher, #A-31570), donkey anti-rabbit Alexa Fluor™ 594 (ThermoFisher, #A-21207), donkey anti-goat Alexa Fluor™ 594 (ThermoFisher, #A-11058), donkey anti-sheep Alexa Fluor™ 647 (ThermoFisher, #A-21448), donkey anti-goat Alexa Fluor™ 647 (ThermoFisher, #A-21448) and donkey anti-rabbit Alexa Fluor™ Plus 647 (ThermoFisher, #A32795TR). Primary and secondary antibodies were diluted in (1X TBS, 1% BSA). After three more washes in TBS, DAPI (Abcam, 1 μg/ml in PBS) was added or not for 5 min at RT in the dark and coverslips were mounted on the slides using Vectashield Antifade mounting medium (Vectorlabs). For thioflavin S staining, a fresh 0.05% Thioflavin S (Sigma-Aldrich) solution was prepared in 100 ml of 50% EtOH and filtered using 0.2 μm filters prior to use. After secondary antibody staining, sections were immersed in thioflavin S solution for 8 min in the dark at RT. Sections were washed two times for 15 seconds with 80% EtOH and then five times in distilled water. Coverslips were mounted on slides as previously described.

For human samples, all steps were carried out in the same fashion to the exception of permeabilization step that lasted 15 min and blocking solution that also contained 0.01% Triton X-100. The following primary and secondary antibodies were used: anti-Lamin B1 antibody (Abcam, #ab16048), anti-DNA (Mouse monoclonal) (Merck, #CBL-186), donkey anti-mouse Alexa Fluor 488 (ThermoFisher, #A-21202) and donkey anti-rabbit Alexa Fluor 555 (ThermoFisher, #A-31572). After secondary antibody incubation, Sudan Black B (Sigma-Aldrich) was incubated for 30 seconds at RT in the dark to quench lipofuscinosis-associated autofluorescence and five washes were performed with distilled water followed by two 5 min washes in PBS. As Sudan black B fluoresces in far-red wavelengths, this channel was not used. For all experiments, all buffers were prepared fresh and filtered with 0.2 μm filters prior to use.

### IHF on free-floating sections

Sections from adult TLR9-GFP mouse brain cut on vibratome were placed in 6-well plate and washed 3 times in PBS. Sections were then incubated with 50 mM NH4Cl for 15 min at RT and then washed two times for 10 min with PBS. Sections were then permeabilized (PBS, 0.25% Triton X-100) for 20 min at room temperature and blocked for 2 h with Blocking solution (1X PBS, 0.25% Triton X-100, 1% BSA). Sections were then incubated for 24 h at 4°C with following primary antibodies: anti-GFP, clones 7.1 and 13.1 (Roche, #11814460001), anti-PGRN antibody (Bio-techne, #AF2557), anti-AIF/Iba1 antibody (Biotechne #NB100-1028) and anti-PLD3 antibody (Proteintech, #17327-1-AP). Sections were then incubated in the dark with following secondary antibodies for 2 h at RT: donkey anti-goat Alexa Fluor™ 488 (ThermoFisher, #A-11055), chicken anti-rabbit Alexa Fluor™ 488 (ThermoFisher, #A-21441), donkey anti-mouse Alexa Fluor™ 555 (ThermoFisher, #A-31570), donkey anti-sheep Alexa Fluor™ 647 (ThermoFisher, #A-21448) and donkey anti-rabbit Alexa Fluor™ Plus 647 (ThermoFisher, #A32795TR). Three washes in PBS were then performed for 10 min each. DAPI (Abcam, 1μg/ml in PBS) was added for 10 min at RT in the dark and sections were washed three times for 10 min with PBS and coaxed on glass slides. Sections were then left to dry and coverslips were mounted using Vectashield Antifade mounting medium (Vectorlabs).

### Image analysis

Plaque quantification: four sections with identical coordinates between genotypes (Bregma -1,655mm, -2,255mm, -2,98mm, -3,455mm) were selected for each 18 months old animal to encompass a large cortical and hippocampal region and stained with anti-Aβ antibody (6E10 clone). Max projections of the 3D Z-stacks were used for analysis on Fiji. To quantify amyloid plaque burden, a threshold function was applied to obtain Aβ plaques and images subjected to the threshold were converted to binary masks. Regions of interest (ROIs) were drawn to delineate hippocampal or cortical regions and their area measured. Amyloid burden was expressed as: Aβ burden = (area of Aβ mask within region)/(total region area). To include the size and the number of plaques, the “Analyze particle” function was used on the binary masks within each region. Corresponding levels with identical coordinates were averaged and compared between genotypes. For 12 months old animals, total number of plaques was manually counted and averaged by animal.

Quantification of plaque-associated PGRN, microglial plaque-associated PGRN and microglia recruitment to the plaques: a total of four sections per animal was used for the analysis. For IBA-1 and PGRN fluorescence intensity, Aβ plaques were converted to an individual ROI after using the "Analyze particle" function on Fiji. The “enlarge” function was used to obtain an expansion of this ROI of 20 μm from the center of each plaque. Integrated density was measured for each individual enlarged ROI. In order to subtract tissue background, two to three measurements in an area with only background signal for the considered channel were drawn, measured and subtracted from the obtained values. Corresponding bregma levels were averaged and compared between genotypes. In order to determine microglial plaque-associated PGRN fluorescence intensity, a minimum of twenty plaques per animal was analyzed. Briefly, on Fiji, the area of PGRN signal was measured within each enlarged ROI (total plaque-associated PGRN area). Following this, IBA-1 binary mask was converted to a selection and PGRN area was measured within this selection in Aβ plaque-enlarged ROIs. Area of microglial plaque-associated PGRN was expressed as a percentage of total plaque-associated PGRN area. To count microglia number near amyloid plaques, twenty individual plaques per animal were selected independently of plaque size. Individual IBA-1 positive cells were manually counted within each Aβ plaque-enlarged ROI and averaged by animal. Segmentation was performed using Ilastik Software.

GPNMB intensity in IBA-1 near amyloid plaques: 3D Z-stack images that include thioflavin S positive Aβ plaques were analyzed from 18 months old APP23 and A9KO mice. IBA-1 channel was converted to a binary mask and then converted to a selection on Fiji. Within Aβ plaque-enlarged ROIs, the integrated density was calculated for GPNMB channel within the IBA-1 selection. Background of the GPNMB channel for each image was subtracted from the obtained values. Segmentation was performed using Ilastik Software.

Quantification of dsDNA: for dsDNA quantification, two sections per animals were stained and four images were acquired for both the hippocampus (CA3 pyramidal layer) and the cortex (entorhinal region) using Leica SP8-gSTED microscope. Max projections for dsDNA, DAPI, PLD3, PGRN and Tom20 were converted to binary masks. On Fiji, to obtain extranuclear dsDNA, the area of dsDNA signal was first measured and the "Image calculator" tool was used to subtract the DAPI mask from the dsDNA mask. The area of the resulting “extranuclear dsDNA” mask was measured and expressed as a percentage of total dsDNA area contained within the field acquired. To quantify the percentage of extranuclear dsDNA inside PLD3, PGRN or Tom20, the “extranuclear dsDNA” mask was multiplied by PLD3 mask to obtain a new mask called “PLD3^+^ extranuclear dsDNA”. The area was measured and expressed as a percentage of total extranuclear dsDNA. The same approach was used for PGRN and Tom20. Segmentation and conversion of signals to binary masks was performed using Ilastik Software. For human scanner acquired sections, the same methodology was applied using Lamin B1 as a proxy for DAPI. Three to four 300 μm x 300 μm images for each region were generated for analysis. Lamin B1 staining was binarized on Fiji using threshold function and area % of nuclear dsDNA was calculated. For extranuclear dsDNA, due to the high variability of signal between patients and healthy controls, area % of extranuclear dsDNA was normalized to total area of the image.

Cell fluorescence intensity on tissue sections: Two to four sections per animals were stained and images were acquired in the hippocampus (CA3 pyramidal layer) and in the cortex (entorhinal region). Brightness contrast was adjusted on max projections on a cytoplasmic marker to better see cell soma in order to delineate cells and draw ROIs for individual cells (only perinuclear cell body and not the dendrites were considered). For each image, background signal was measured in order to subtract it from the measurements. Mean grey values and integrated density for each cell in the considered channel was measured and CTCF was calculated as previously described using the following formula: CTCF = (integrated density - (area of selected cell x mean fluorescence of background readings).

Analysis of PLD3^+^ and PGRN^+^ cells: Analysis on 12 months old animals was done on QuPath software. Four to five sections were analyzed by animal. Briefly, in a first step, ROIs for the hippocampus and the cortex were created. Several identical square annotation regions were created in these regions to generate a sparse image allowing to take into account heterogeneity in background and signal intensity on all sections analyzed. Stardist training for detection of nuclei (minimum nuclei area 5, probability detection threshold of 0.5, with the options “include probability”, “cell expansion 5” and “constrain cell expansion”) was carried out on the sparse image. For non-microglial cell analysis, nuclei belonging to IBA-1 positive cells were excluded (“Ignore*” annotation). For analysis on microglia, only nuclei that belonged to IBA-1^+^cells were selected during the training. After nuclei detection, new object classifiers (random trees) were created and trained on the sparse image first for PGRN and then for PLD3 channel. Only cells that had intense puncta in the perinuclear area were considered as positive. Following this, a composite object classifier was created and batched on all images in the region considered and verified image by image. Proportions of positive cells in percentages were calculated from exported data using the number of PLD3^+^, or PGRN^+^ cells from the total number of nuclei detected in the region considered.

Neuronal loss in CA1 layer: pyramidal layer thickness and nuclei density analysis in CA1 region was performed on 18 months old animals using scanner-acquired images of the left hemisphere. Three images belonging to three corresponding different bregma level for each animal were analyzed. New images were created to encompass 1 mm of CA1 pyramidal layer. To analyze the thickness of the layer on Fiji, three measurement were made throughout the layer using DAPI channel and averaged by image. Averaged thickness measurements were then averaged by animal. For the number of nuclei within the layer, Cellpose plugin with cyto2 model (DAPI: nuclei channel, PGRN: cytoplasmic channel, 20-25μm diameter threshold depending on the image) was used to segment the nuclei and obtain binary masks. A ROI was then created to encompass cells belonging to the pyramidal layer. After correcting masks (“Watershed” function) to separate individual nuclei that clustered during segmentation, the “analyze particle” function was used to obtain the number of cells within the ROI layer excluding very small nuclei (<5 μm) and particles on the edge of the image. Number of cells were averaged by animal.

### Behavioral tests

All experiments were carried out in the same facility. Before testing, animals were transferred to the behavior animal house (IBPS, Jussieu, Paris) and left two weeks unperturbed for acclimatation. Handling with the experimenter was carried out each day for one week prior to behavioral testing. All experiments were performed in a quiet environment. Before each test, animals were isolated in individual cages for at least 1 h in the experimental room. The order of passage of the animals for each test was random and apparatuses were cleaned between two animals first with water then with 20% EtOH.

Novel Object Recognition: four 40 cm x 40 cm boxes were placed in the room and the light intensity was set uniformly at 8-10 lux in all the boxes for the duration of the test. The test was carried out on five consecutive days consisting of four sessions of habituation and two test sessions. For each session, animals were placed in the box with the muzzle facing the same corner for each individual box. The first three habituation sessions, animals were left to explore the empty box for 20 min. Test sessions only lasted 10 min. On the fourth habituation session, two identical objects were placed in the box (aligned, 10 cm from the corners). One hour later, one of the identical objects (familiar objects: FO) was placed to the opposite side of the box. The next day, one of the identical objects was replaced with a new object (NO; same positions than for the first habituation session with identical objects). All trials were recorded using a camera and Ethovision XT software. Total distance travelled and velocity were recorded by the software. For object exploration analysis, all trials were manually timed considering that the animals are exploring the object when the muzzle is close and facing the object and/or if the front paws are resting on the object. Animals that spent less then 5 seconds exploring each object during the 10 min of testing period were excluded from the analysis. Total exploration time for each object was timed and discrimination ratio (DR) was expressed as:

DR = (exploration time NO – exploration time FO) / (total time of object exploration).

Y-Maze: a three-arm maze with 120° angle between arms and different visual cues on the walls of each arm was used. For testing, light was set at 100 lux uniformly in the Y-Maze. Individual animals were placed in the three-arm maze, left to explore for 5 min and tracked using Imetronic software. Animals were placed in the Y-Maze always in the same arm and in the same direction. For analysis, total number of arm entries was counted and tracking of the animal allowed for recording of each animal’s itinerary. The percentage of alternations was then expressed as:

% of alternation = (total number of alternations) / (total number of arm entries - 2).

Open Field: four 40 cm x 40 cm boxes were placed in the room and the light intensity was set uniformly at 150 lux in all the boxes for the duration of the test. Using Ethovision XT software, 30 cm x 30 cm square detection regions in the center of each box were created. Animals were placed in the box and left to explore for 20 min. Total distance travelled, velocity and time spent in the center were recorded by the camera/software and analyzed.

Light-dark box: four 20 cm x 40 cm boxes with a lit chamber and a dark chamber (20 cm x 30 cm for lit chamber and 20 cm x 10 cm for dark chamber) were placed in the room and the light intensity was uniformly set at 300 lux in in the lit chamber. Using Ethovision XT software, detection regions encompassing the lit chamber were created. Animals were placed in the dark chamber of the box and left to explore for 10 min. Total distance travelled, velocity and time spent in the lit chamber were recorded by the camera and software and analyzed.

Tail suspension test: a camera was used to record animals and recording was started before placing the first animal. Five animals were tested per individual session. Animals were suspended using strongly adherent tape on their tails and left for 5 min. Using the recordings, immobility time was manually timed for each animal exactly 5 min from the start of the suspension.

### Statistical analysis

Statistics were calculated with GraphPad Prism software version 9.0 (GraphPad Inc.) and p-values were determined using either two-tailed unpaired t-tests or one-way ANOVA with Tukey’s multiple comparison test, consistently comparing all groups with each other. Two-tailed t-tests were used for comparison between two groups and one-way ANOVA when comparing more than two groups. All graphs show mean ± standard error of mean (SEM). Information regarding replicates can be found in the corresponding figure legend.

## List of reagents used

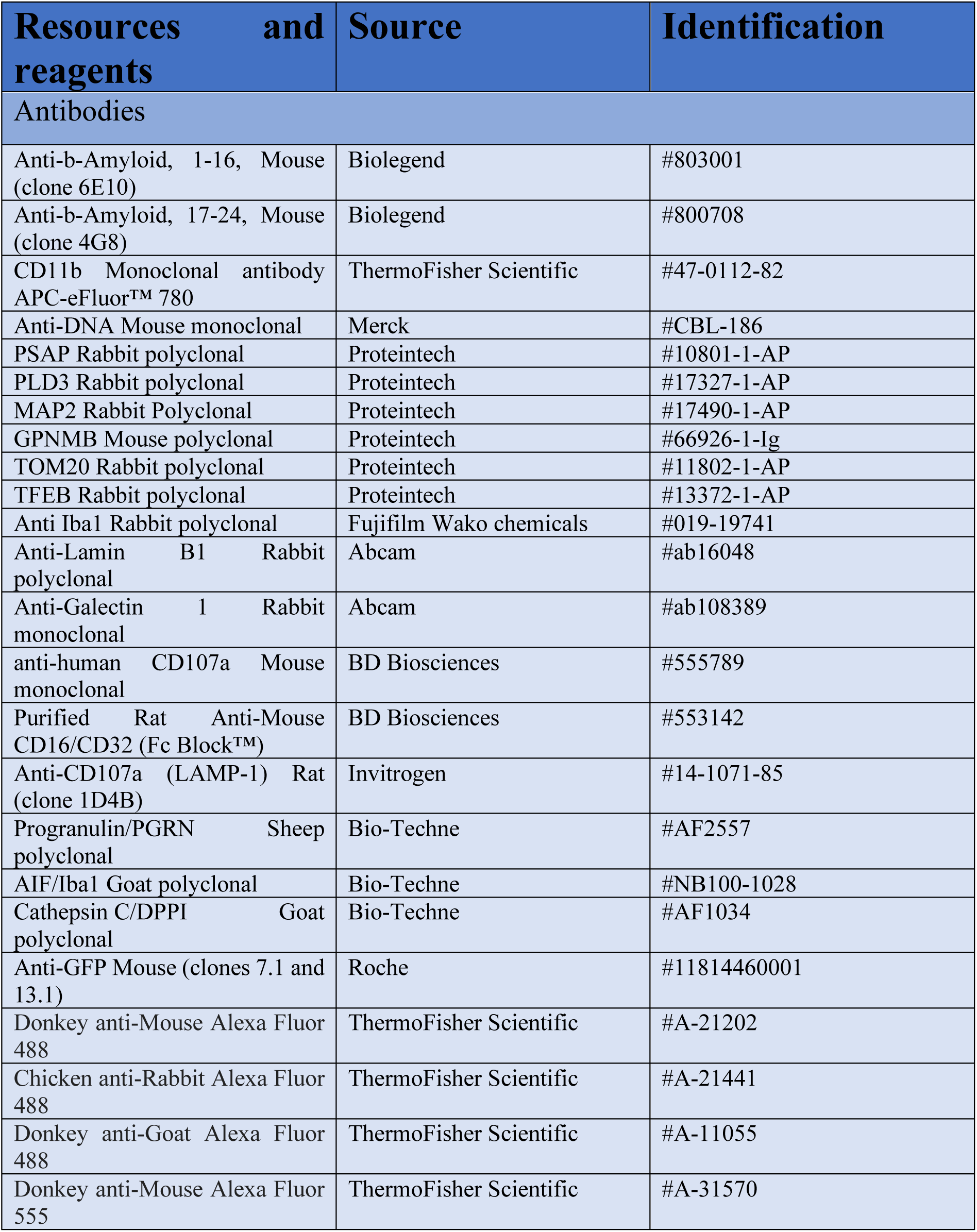
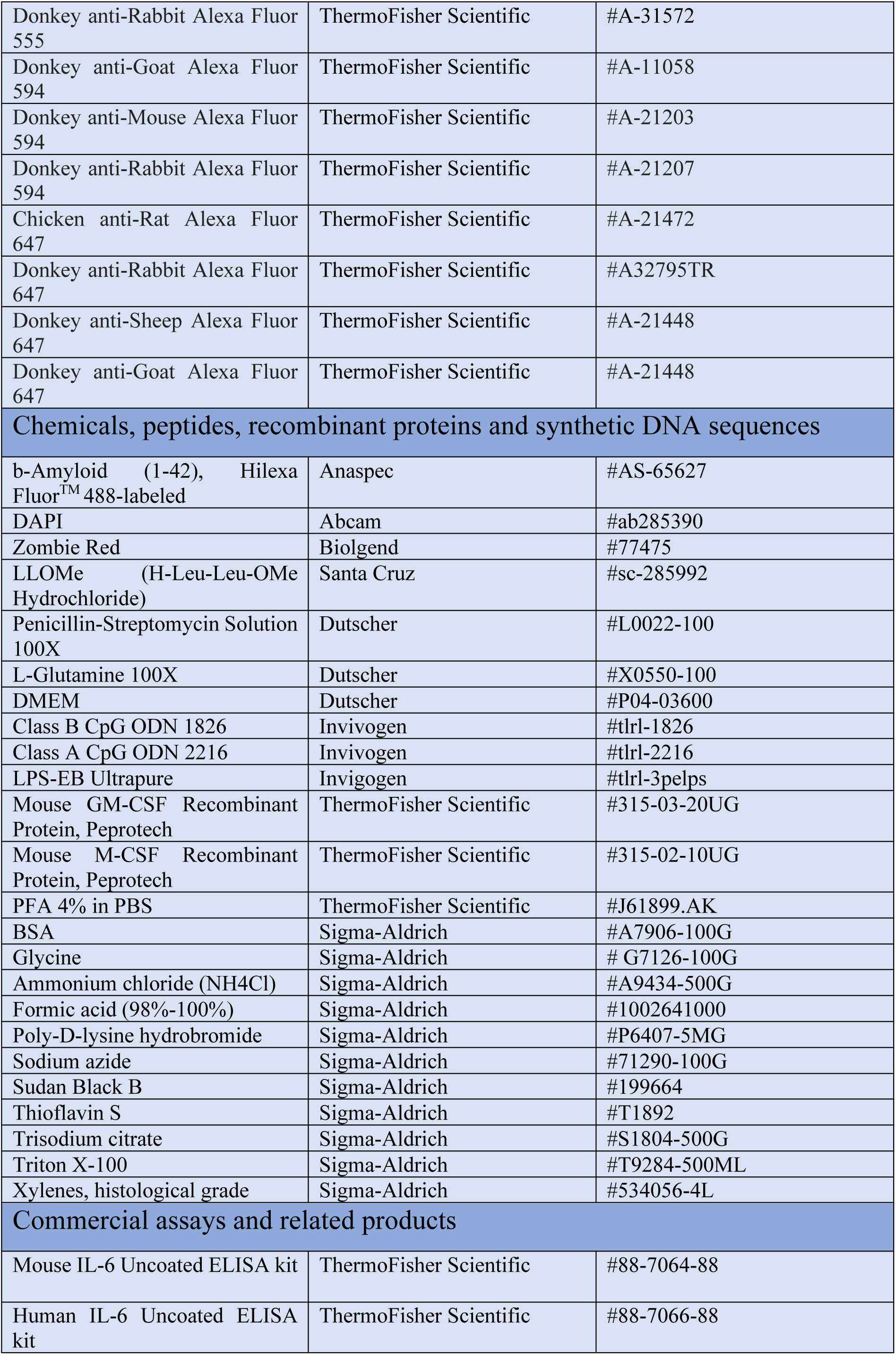
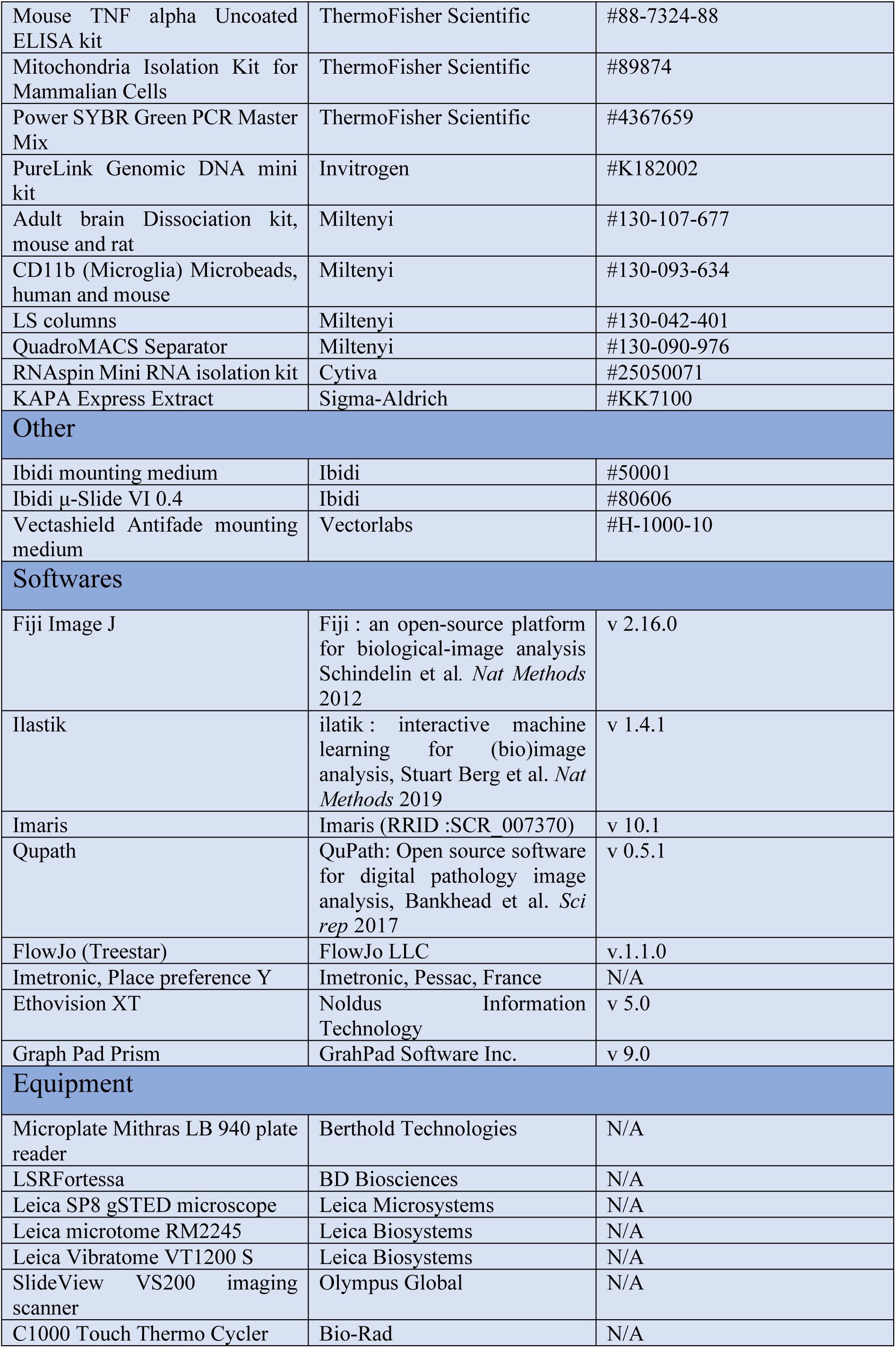

## Data availability

The data are available upon request from

## Acknowledgements

This study as supported by grants from Fondation Vaincre Alzheimer, Fondation Alzheimer and INSERM; an MRT and ALZProtect fellowships to MDL and an MRT fellowship to LM. The authors are grateful to Dr S. Akira for providing the TLR9-KO mice; to Pr. M. Brinkmann for providing TLR9-GFP mice; to Nicolas Goudin (INEM) for his assistance regarding image analysis and macro design; to Meriem Garfa-Traore from the SFR Necker Cellular Imagery platform; to Sophie Berissi and all the staff from the SFR Necker Histology platform and Histomics platform of the ICM.

## Author contributions

BM and MDL designed the study and LM, KP, MDL, IE, GDJ and BM performed experiments, analyzed and interpreted data. MCP, SV, CE and PV provided critical research tools. BM and MDL wrote the manuscript and BM supervised the whole project.

