## Supplementary Figures for "Toll-like receptor 9 contributes in microglial activation and lysosomal dysfunction to promote Alzheimer’s disease"

### Figure S1

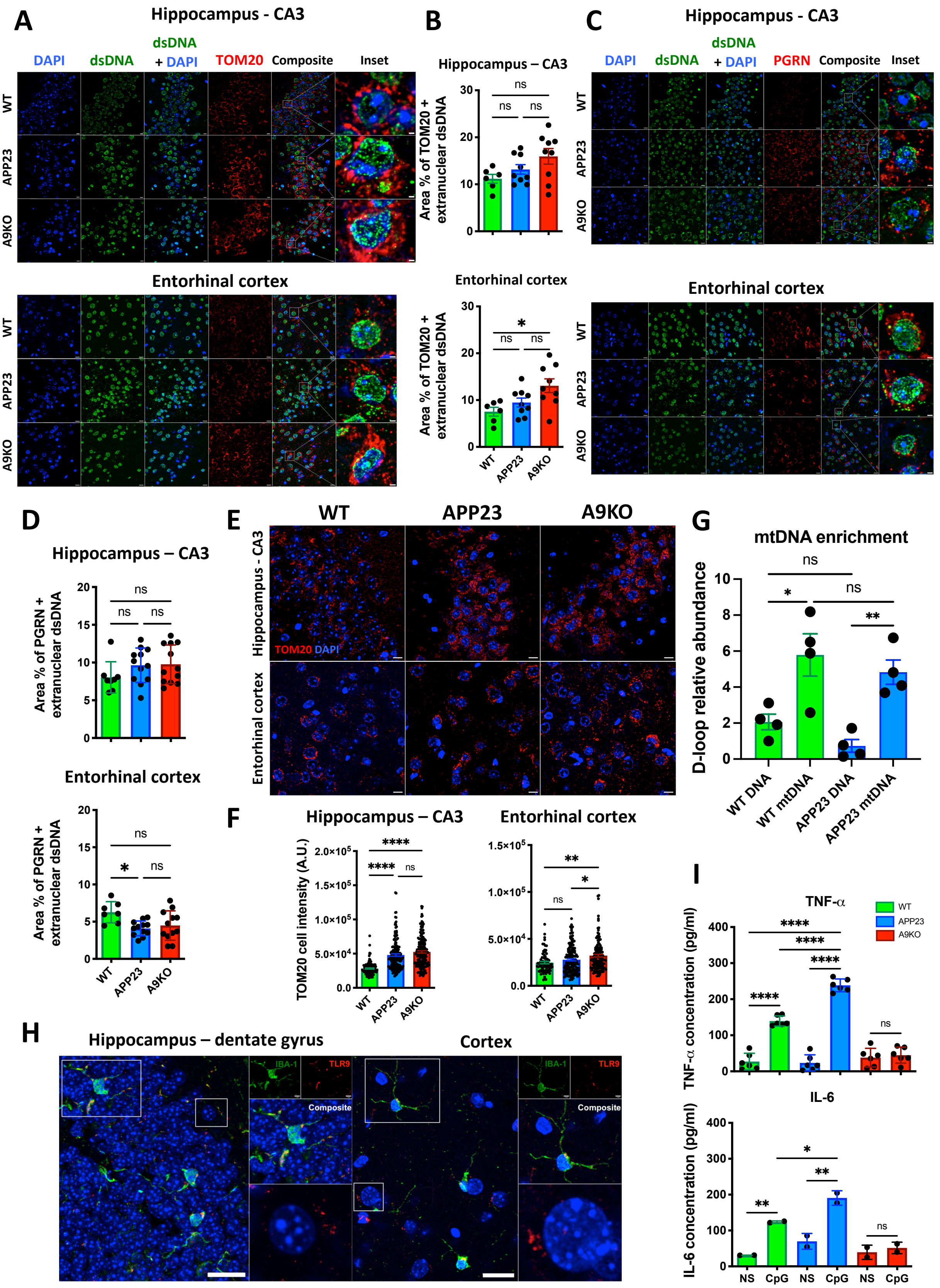

A
Hippocampus - CA3
DAPI
dsDNA
dsDNA + DAPI
TOM20
Composite
Inset
WT
APP23
A9KO
Entorhinal cortex
B
Hippocampus – CA3
Area % of TOM20 + extranuclear dsDNA
Entorhinal cortex
C
PGRN
D
Area % of PGRN + extranuclear dsDNA
E
TOM20 DAPI
Hippocampus - CA3
F
TOM20 cell intensity (A.U.)
G
mtDNA enrichment
D-loop relative abundance
WT DNA
WT mtDNA
APP23 DNA
APP23 mtDNA
H
Hippocampus – dentate gyrus
Cortex
IBA-1
TLR9
Composite
I
TNF-α
TNF-α concentration (pg/ml)
IL-6
IL-6 concentration (pg/ml)
NS
CpG
ns

### Figure S2

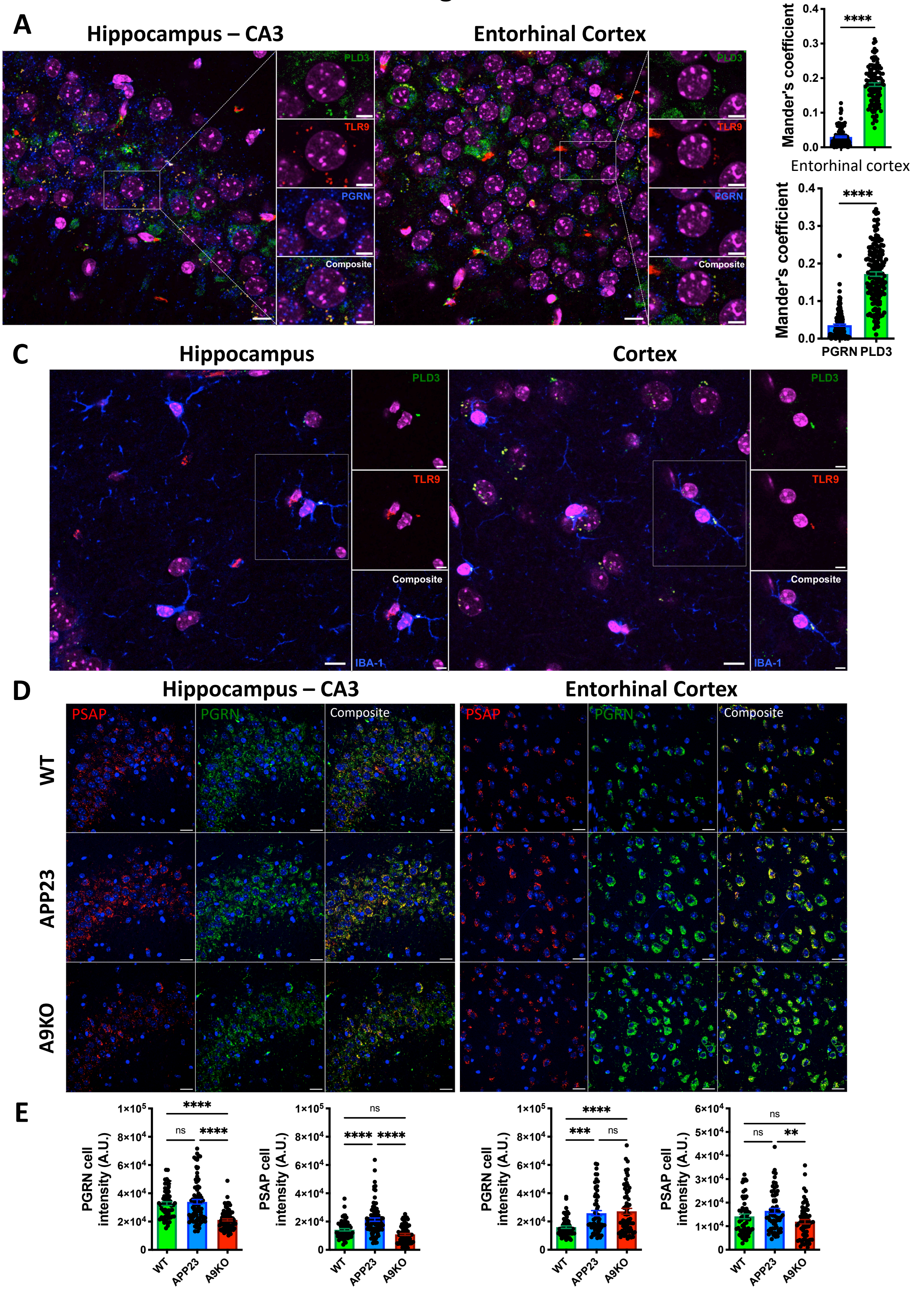

A
Hippocampus – CA3
Entorhinal Cortex
PLD3
TLR9
PGRN
Composite
B
Hippocampus – CA3
Mander's coefficient
****
Entorhinal cortex
PGRN
PLD3
C
Hippocampus
Cortex
IBA-1
D
PSAP
WT
APP23
A9KO
E
PGRN cell intensity (A.U.)
PSAP cell intensity (A.U.)
ns
***
**

### Figure S3

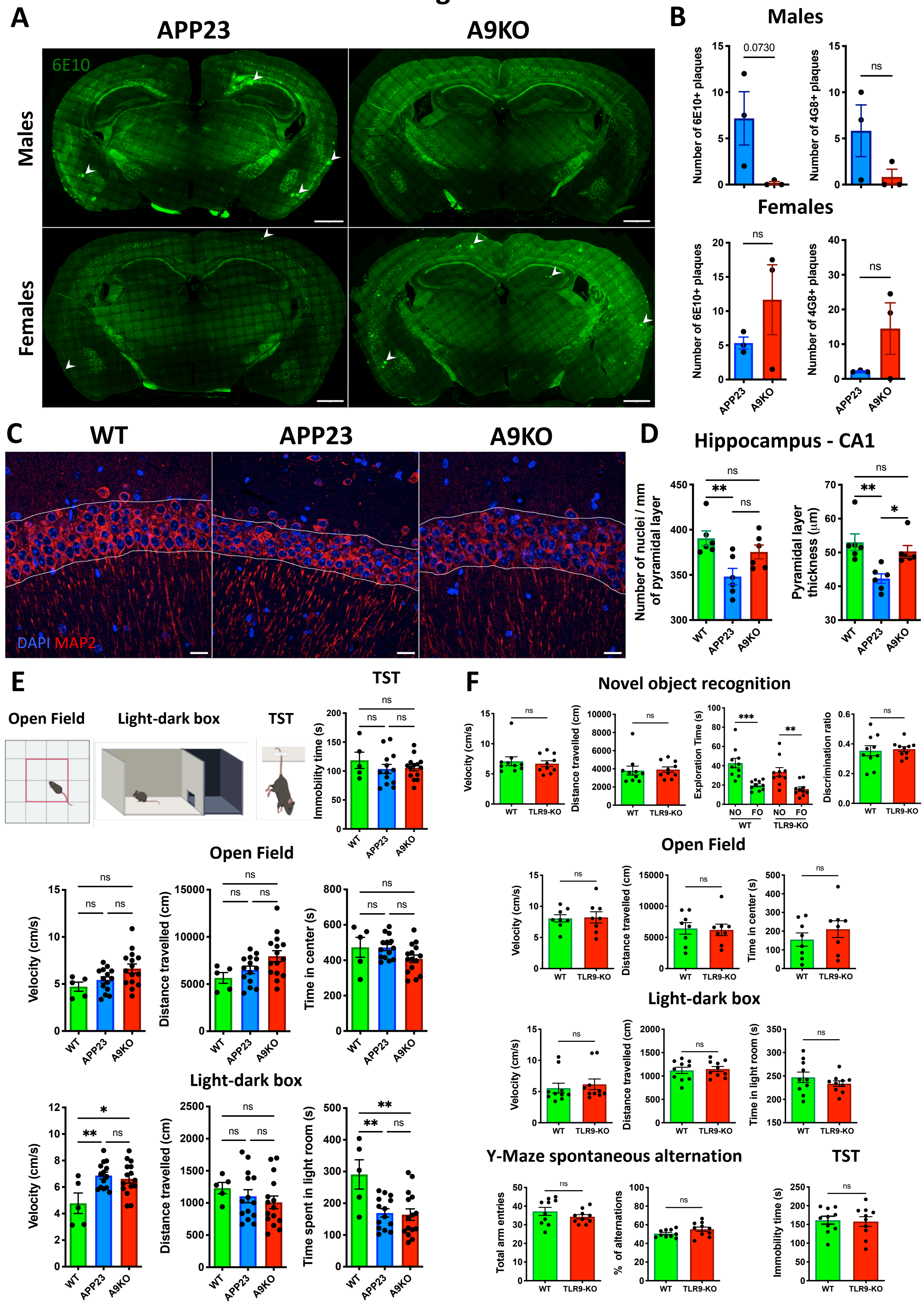
A
APP23
A9KO
Males
Females
6E10
B
Males
Number of 6E10+ plaques
0.0730
Number of 4G8+ plaques
ns
Females
APP23
A9KO
C
WT
APP23
A9KO
DAPI MAP2
D
Hippocampus - CA1
Number of nuclei / mm of pyramidal layer
Pyramidal layer thickness (µm)
E
Open Field
Light-dark box
TST
Immobility time (s)
Velocity (cm/s)
Distance travelled (cm)
Time in center (s)
Time spent in light room (s)
F
Novel object recognition
Exploration Time (s)
Discrimination ratio
NO
FO
TLR9-KO
Time in light room (s)
Y-Maze spontaneous alternation
Total arm entries
% of alternations

### Figure S4

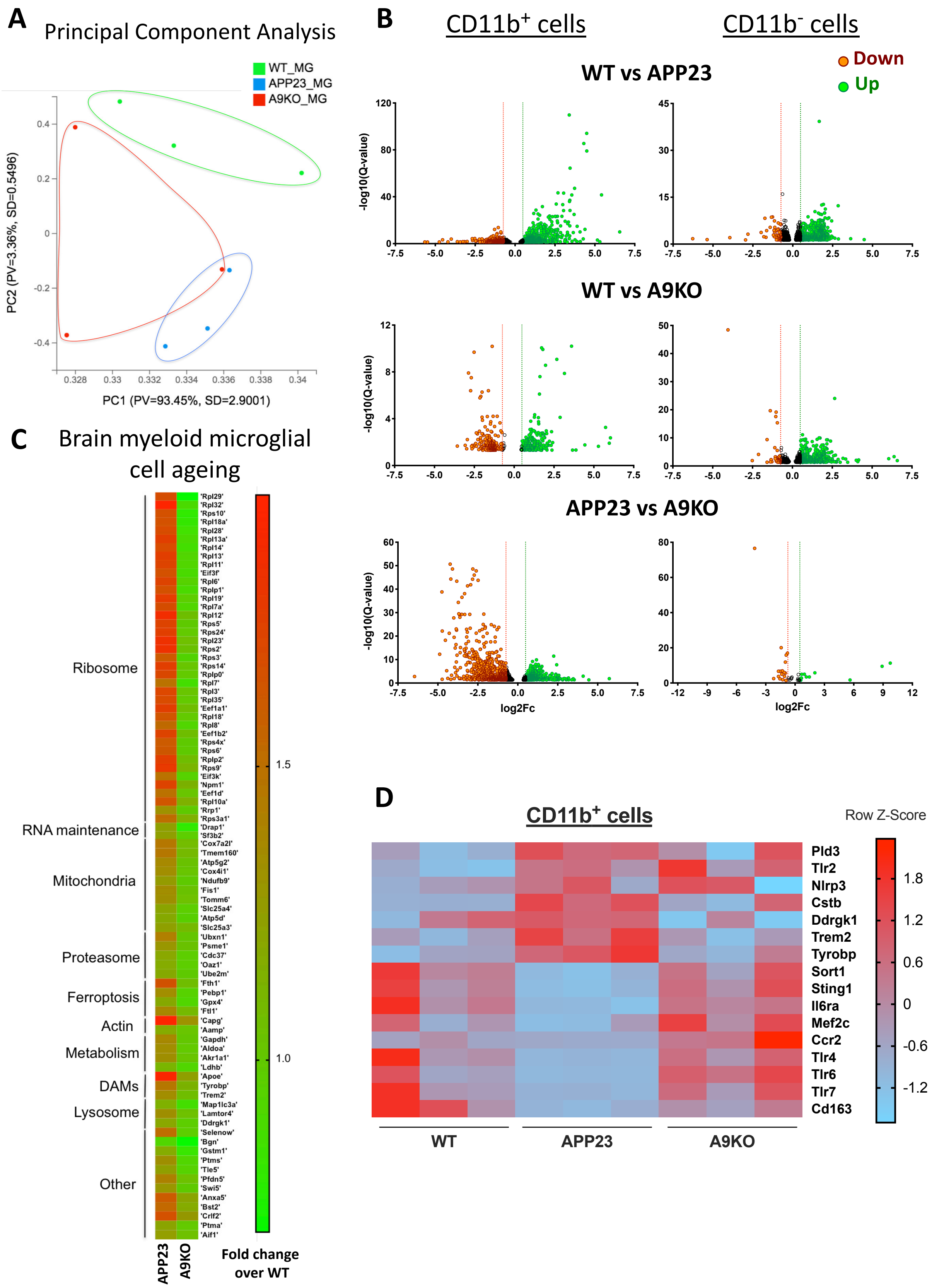

A
Principal Component Analysis
WT_MG
APP23_MG
A9KO_MG
PC2 (PV=3.36%, SD=0.5496)
PC1 (PV=93.45%, SD=2.9001)
B
CD11b+ cells
CD11b- cells
Down
Up
WT vs APP23
WT vs A9KO
APP23 vs A9KO
-log10(Q-value)
log2Fc
C
Brain myeloid microglial cell ageing
Ribosome
RNA maintenance
Mitochondria
Proteasome
Ferroptosis
Actin
Metabolism
DAMs
Lysosome
Other
'Rpl29' 'Rpl32' 'Rps10' 'Rpl18a' 'Rpl28' 'Rpl13a' 'Rpl14' 'Rpl13' 'Rpl11' 'Eif3f' 'Rpl6' 'Rplp1' 'Rpl19' 'Rpl7a' 'Rpl12' 'Rps5' 'Rps24' 'Rpl23' 'Rps2' 'Rps3' 'Rps14' 'Rplp0' 'Rpl7' 'Rpl3' 'Rpl35' 'Eef1a1' 'Rpl18' 'Rpl8' 'Eef1b2' 'Rps4x' 'Rps6' 'Rplp2' 'Rps9' 'Eif3k' 'Npm1' 'Eef1d' 'Rpl10a' 'Rrp1' 'Rps3a1' 'Drap1' 'Sf3b2' 'Cox7a2l' 'Tmem160' 'Atp5g2' 'Cox4i1' 'Ndufb9' 'Fis1' 'Tomm6' 'Slc25a4' 'Atp5d' 'Slc25a3' 'Ubxn1' 'Psme1' 'Cdc37' 'Oaz1' 'Ube2m' 'Fth1' 'Pebp1' 'Gpx4' 'Ftl1' 'Capg' 'Aamp' 'Gapdh' 'Aldoa' 'Akr1a1' 'Ldhb' 'Apoe' 'Tyrobp' 'Trem2' 'Map1lc3a' 'Lamtor4' 'Ddrgk1' 'Selenow' 'Bgn' 'Gstm1' 'Ptms' 'Tle5' 'Pfdn5' 'Swi5' 'Anxa5' 'Bst2' 'Crlf2' 'Ptma' 'Aif1'
APP23
A9KO
Fold change over WT
1.5
1.0
D
CD11b+ cells
Row Z-Score
Pld3
Tlr2
Nlrp3
Cstb
Ddrgk1
Trem2
Tyrobp
Sort1
Sting1
Il6ra
Mef2c
Ccr2
Tlr4
Tlr6
Tlr7
Cd163
1.8
1.2
0.6
0
-0.6
-1.2
WT
APP23
A9KO

### Figure S5

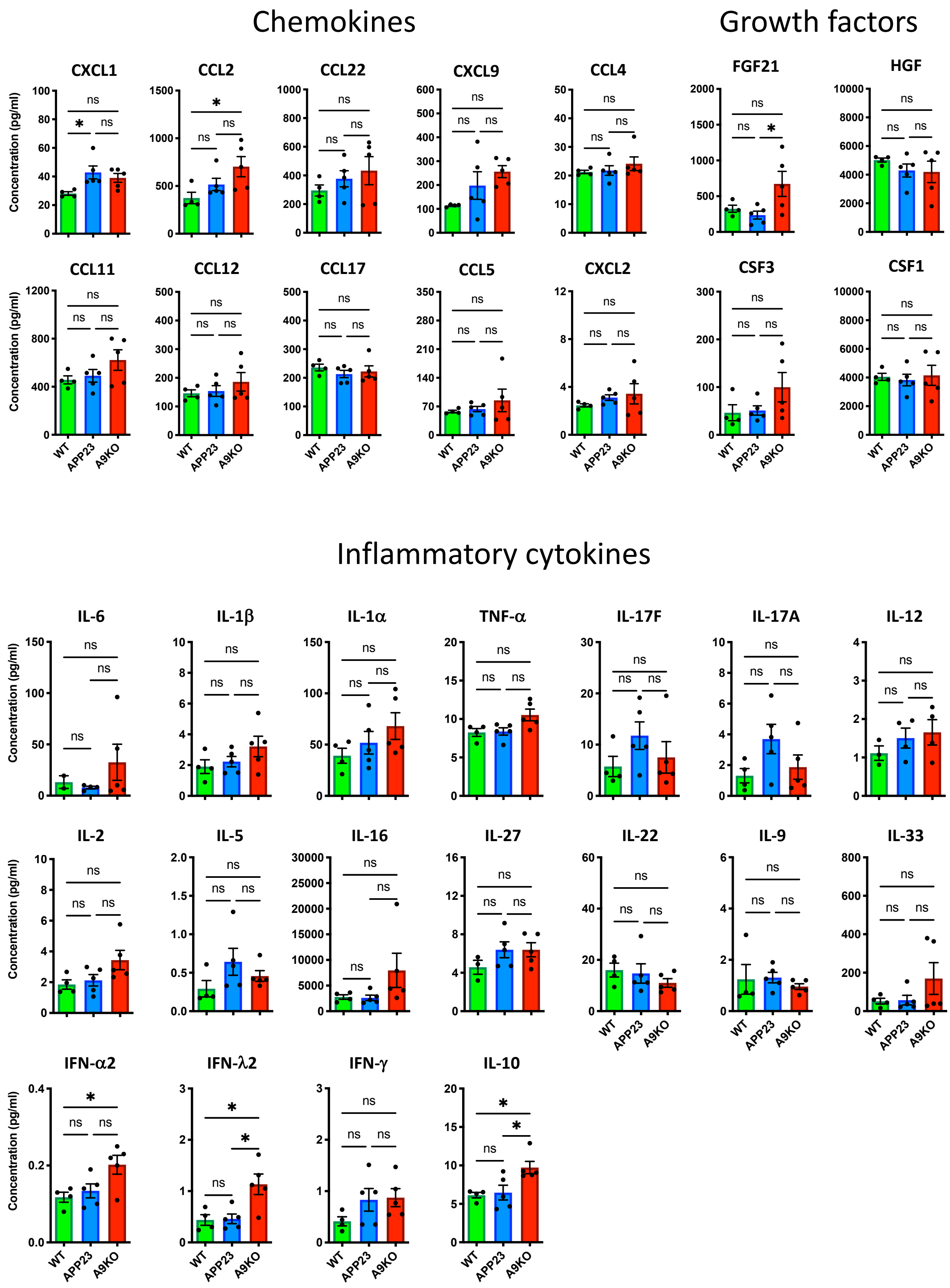
Chemokines
Growth factors
CXCL1
CCL2
CCL22
CXCL9
CCL4
FGF21
HGF
CCL11
CCL12
CCL17
CCL5
CXCL2
CSF3
CSF1
Inflammatory cytokines
IL-6
IL-1β
IL-1α
TNF-α
IL-17F
IL-17A
IL-12
IL-2
IL-5
IL-16
IL-27
IL-22
IL-9
IL-33
IFN-α2
IFN-λ2
IFN-γ
IL-10
Concentration (pg/ml)
WT
APP23
A9KO
ns
*

### Figure S6

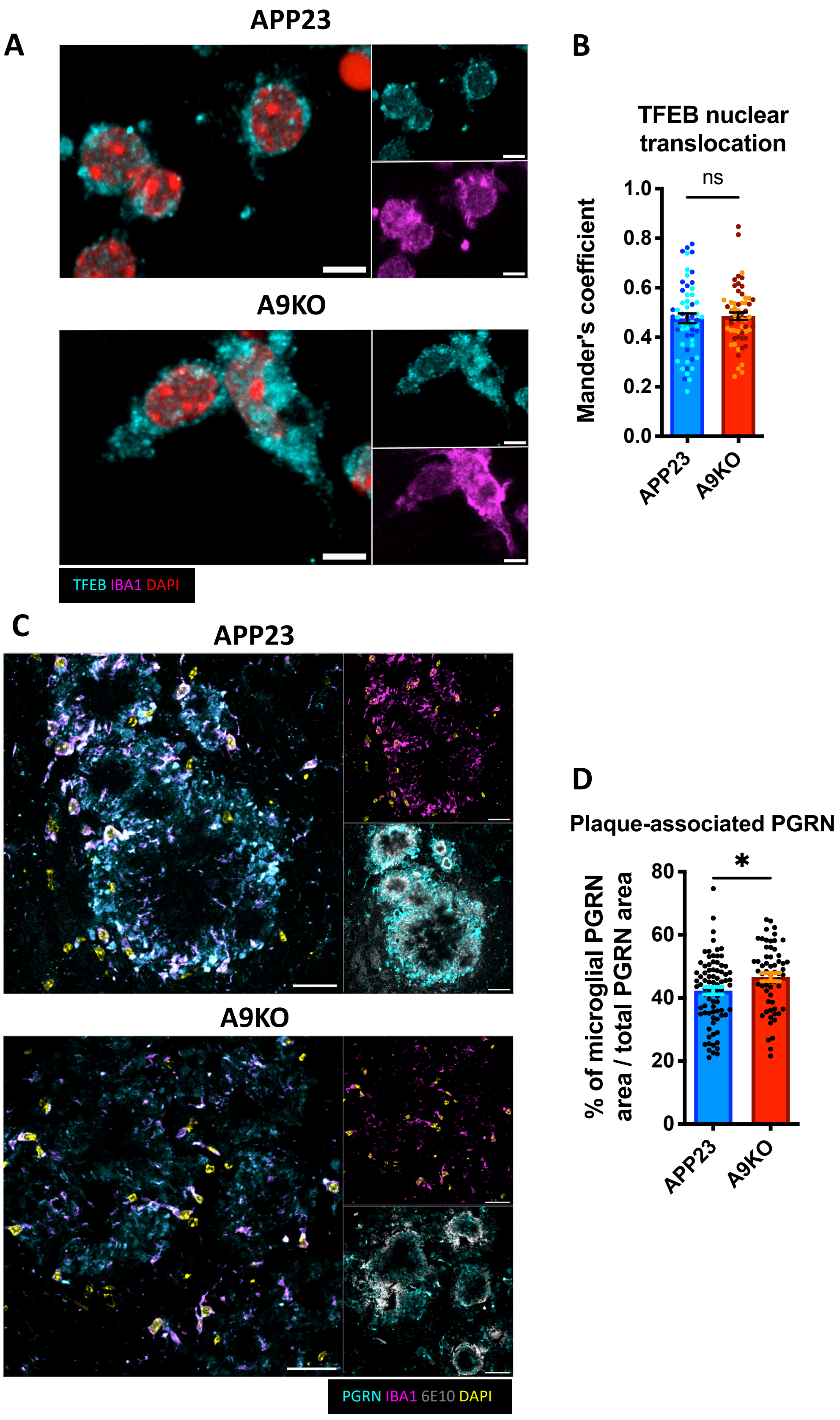

A
APP23
A9KO
TFEB IBA1 DAPI
B
TFEB nuclear translocation
ns
Mander's coefficient
1.0
0.8
0.6
0.4
0.2
0.0
APP23
A9KO
C
APP23
A9KO
PGRN IBA1 6E10 DAPI
D
Plaque-associated PGRN
*
% of microglial PGRN area / total PGRN area
80
60
40
20
0
APP23
A9KO

**Figure S1. dsDNA localization and TLR9-mediated microglia activation**

**(A)** Immunofluorescence images of 12 months old WT (n=2), APP23 (n=3) and A9KO (n=3) mice stained for dsDNA (green), mitochondria (TOM20, red) and nucleus (DAPI, blue) in the CA3 region of the hippocampus and in the entorhinal cortex. Scale bar: 10 μm. Scale bar magnification: 2 μm.

**(B)** Quantification $TOM20^+$ extranuclear dsDNA. Each dot represents one image quantified (n=3 images per animal). n=2-3. ns: not significative, *p<0.05; one-way ANOVA was performed with Tukey post-hoc test.

**(C)** Immunofluorescence images of 12 months old WT (n=2), APP23 (n=3) and A9KO (n=3) mice stained for dsDNA (green), PGRN (red) and nucleus (DAPI, blue) in the CA3 region of the hippocampus and in the entorhinal cortex. Scale bar: 10 μm. Scale bar magnification: 2 μm.

**(D)** Quantification of $PGRN^+$ extranuclear dsDNA. Each dot represents one image (n=3-4 images per animal). n=2-3. ns: not significative, *p<0.05; one-way ANOVA was performed with Tukey post-hoc test.

**(E)** Immunofluorescence images of 12 months old WT (n=2), APP23 (n=3) and A9KO (n=3) mice stained for mitochondria (TOM20, red) and nucleus (DAPI, blue) in the CA3 region of the hippocampus and in the entorhinal cortex. Scale bar: 15 μm.

**(F)** TOM20 relative cell fluorescence intensity. n=20 cells or more per section was analyzed (n=3 sections per animal). n=2-3. ns: not significative, *p<0.05, **p<0.01, ****p<0.0001; one-way ANOVA was performed with Tukey post-hoc test.

**(G)** Murine mitochondrial DNA from whole brain (mtDNA) and genomic DNA (DNA) from tail of 15 months old WT and APP23 mice analyzed by qPCR for mitochondrial D-loop. Each point represents relative abundance of D-loop normalized to *Actb* (β-actin) gene for one dilution of the sample. ns: not significative, *p<0.05, **p<0.01; one-way ANOVA was performed with Tukey post-hoc test.

**(H)** Immunofluorescence images of TLR9-GFP mice (n=3) stained for TLR9 (anti-GFP, red), microglia (IBA-1, green) and nucleus (DAPI, blue) in the dentate gyrus region of the hippocampus and in the cortex. Scale bar composite: 20 μm. Scale bar frame: 5 μm.

**(I)** IL-6 and TNF-α production by microglia purified from 12 to 15 months old WT, APP23 and A9KO mice, stimulated with 1 μg/ml of CpG-B and measured by ELISA. n=3 for TNF-α; n=1 for IL-6. ns: not significative, *p<0.05, **p<0.01, ****p<0.0001; one-way ANOVA was performed with Tukey post-hoc test. NS: non-stimulated.

**Figure S2. TLR9 is associated with lysosomal PLD3 and modulates PGRN and PSAP expression**

**(A)** Immunofluorescence images of TLR9-GFP mice (n=3) stained for TLR9 (anti-GFP, red), PLD3 (green), PGRN (blue) and nucleus (DAPI, magenta) in the CA3 region of the hippocampus and in the entorhinal cortex. Scale bar: 10 µm. Scale bar magnification: 5 µm.

**(B)** Colocalization of TLR9 with PGRN or PLD3 expressed as Mander's coefficient. n=114 cells for CA3 region and n=166 cells for entorhinal cortex. ****p<0.0001; two-tailed student's t-test was performed.

**(C)** Immunofluorescence images of TLR9-GFP mice (n=3) stained for TLR9 (anti-GFP, red), PLD3 (green), microglia (IBA-1, blue) and nucleus (DAPI, magenta) in the CA3 region of the hippocampus and in the cortex. Scale bar: 10 µm. Scale bar individual cell: 5 µm.

**(D)** Immunofluorescence images of 12 months old WT (n=2), APP23 (n=3) and A9KO (n=3) mice stained for PSAP (red), PGRN (green) and nucleus (DAPI, blue) in the CA3 region of the hippocampus and in the entorhinal cortex. Scale bar: 20 µm. PSAP: prosaposin**.**

**(E)** PGRN (left) and PSAP (right) relative cell fluorescence intensity. CA3: WT=58 cells, APP23=86 cells, A9KO=83 cells; entorhinal cortex: WT=59 cells, APP23=64 cells, A9KO=65 cells. ns: not significative, **p<0.01, ***p<0.001, ****p<0.0001, one-way ANOVA was performed with Tukey post-hoc test.

**Figure S3. Early Aβ deposition, neuronal loss and cognitive performance in APP23 and A9KO mice**

**(A)** Immunofluorescence images of whole brain sections from 12 months old mice (n=3 for males and females) stained for Aβ plaques (6E10, green). Arrows point to individual plaques. Scale bar: 1 mm.

**(B)** Total number of plaques for males (top) and females (bottom). n=2 sections per animal for both 6E10 and 4G8 were analyzed. Each dot corresponds to the average number of plaques for one animal. ns: not significative; two-tailed student's t-test was performed.

**(C)** Immunofluorescence images of 18 months old WT (n=6), APP23 (n=6), A9KO (n=6) mice stained for neurons (MAP-2, red) and nucleus (DAPI, blue) in the CA1 pyramidal layer of the hippocampus. The edge of the layer is manually delineated. Scale bar: 20 µm.

**(D)** Number of nuclei and pyramidal layer thickness in the CA1 region (1 mm of layer). n=3 sections per animal in different bregma levels were analyzed. Each dot represents one animal. ns: not significative, *p<0.05, **p<0.01; one-way ANOVA was performed with Tukey post-hoc test.

**(E)** Additional behavioral tests performed in 12 months old animals: open field for anxiety and locomotion (WT n=5, APP23 n=14, A9KO n=15), tail suspension (TST) for depression (WT n=5, APP23 n=13, A9KO n=14) and light-dark box for anxiety (WT n=5, APP23 n=14, A9KO n=15). TST: Immobility time (top right); open field: velocity, distance travelled and time spent in the center of the arena (middle); light-dark box: velocity, distance travelled and time spent in the light room (bottom). ns: not significative, *p<0.05, **p<0.01; one-way ANOVA was performed with Tukey post-hoc test.

**(F)** Behavioral tests performed in healthy 9 months old WT and TLR9-KO mice. NOR (top), open field and light-dark box (middle), Y-Maze (bottom left) and TST (bottom right). n=10 for all tests except for the open field (n=8). ns: not significative, **p<0.01, ***p<0.001; two-tailed student's t-test was performed. Concerning the NOR, NO exploration time was compared to FO exploration time for the same genotype. NO: Novel Object, FO: Familiar Object.

**Figure S4. TLR9 expression drives CNS-resident cells transcriptional signature**

**(A)** Principal component analysis on purified microglia from 18 months old WT, APP23 and A9KO mice. n=3. PV: Proportion of variance; PC: Principal component; SD: Standard deviation.

**(B)** Volcano plots showing DEGs between WT and APP23 (top), WT and A9KO (middle), APP23 and A9KO (bottom) either in $CD11b^+$ or in $CD11b^-$ cells. Differential expression was assessed with -log10(Q-Value) and log2Fc. Threshold bars show log2Fc cut-off used for differential expression analysis.

**(C)** Heatmap of the 84 significant genes (Q-value<0.05) from all genes found in the "myeloid and microglial cell ageing" gene-set from the GSEA analysis for APP23 and A9KO compared to WT microglia. DAMs: disease-associated microglia. Heatmap shows mean (per genotype) fold change over WT microglia.

**(D)** Transcriptional analysis of additional DEGs in WT, APP23 and A9KO microglia. Heatmap shows row Z-Score. Only DEGs that passed the Q-value cut-off are shown here except for NLRP3 that is displayed to illustrate no significant difference.

**Figure S5. TLR9 induces small changes in inflammatory mediators at the periphery during Aβ deposition**

Concentration of chemokines, growth factors and inflammatory cytokines in the serum of 12 months old animals (WT n=4, APP23 n=5, A9KO n=5). Each dot represents one animal. ns: not significative, *$p<0.05$; one-way ANOVA was performed with Tukey post-hoc test.

**Figure S6. TLR9 contributes to lysosomal fitness in microglia**

**(A)** Immunofluorescence images of purified microglia from 10 to 15 months old APP23 and A9KO mice stained for TFEB (cyan), microglia (IBA-1, magenta) and nucleus (DAPI, red). n=2. Scale bar: 5 µm.

**(B)** Colocalization of TFEB with DAPI expressed as Mander's coefficient. n= 20 cells or more per animal were analyzed. n=2. ns: not significative; two-tailed student's t-test was performed.

**(C)** Immunofluorescence images of 18 months old APP23 and A9KO mice stained for Aβ plaques (6E10, grey), PGRN (cyan), microglia (IBA-1, magenta) and nucleus (DAPI, yellow) in the cortex. n=3. Scale bar: 20 µm.

**(D)** Quantification of microglial-PGRN area / total PGRN area in Aβ plaque vicinity. n=81 plaques for APP23 and n=61 plaques for A9KO mice. Each dot represents an individual plaque. n=3. *$p<0.05$; two-tailed student's t-test was performed.
